# A dynamical circuit model of *C. elegans* chemotaxis with emergent sharp turns

**DOI:** 10.64898/2026.08.09.743732

**Authors:** Ada Squires, Victoria Booth, Eleni Gourgou

## Abstract

With its rigorously characterized connectome, the nematode *Caenorhabditis elegans* is a powerful model organism to study the fundamental roles of neuronal circuits in behavior. However, despite the breadth of research, many questions remain unanswered regarding how these organisms are able to successfully navigate their environment. Here, we present a biologically grounded dynamical circuit model for the investigation of sensory-guided behavior during *C. elegans* chemotaxis. Our model consists of the chemosensory neuron AWA, interneurons RIM and RIA, motor neurons, including SMDs and RMDs, and body wall muscles that provide proprioceptive feedback through stretch receptors. The connectivity prioritizes functional and dynamical correspondence with the biological circuit over one-to-one anatomical fidelity. After optimization with an evolutionary algorithm, the model locomotes toward a chemical attractant, effectively capturing nematode chemotactic behavior. A key emergent property of the model that contributes to successful chemotaxis is the ability to make sharp turns, which resemble the omega turns of living nematodes. The direction and magnitude of these turns depend on the phase of the ongoing locomotory cycle. The sharp turning behavior is triggered by decreases in the concentration of the attractant. In the model, decreases in attractant concentration levels reduce AWA activity, which in turn triggers disinhibition of RIM activity and subsequent changes in RIA oscillatory activity. The ensuing coordinated changes in downstream motor neurons’ activity patterns produce sharp turns, which correct the model worm’s path to head toward the attractant source and, after reaching the concentration gradient peak, allow the model worm to remain in its proximity. The proposed framework and its emergent dynamics provide new insights into how circuit-level dynamics may generate key features of *C. elegans* chemotactic behavior, including omega-like turns. In parallel, it generates experimentally testable hypotheses about how the participating neuronal elements contribute to chemotactic behavior.

## 1. Introduction

With only 302 neurons and a wide repertoire of behaviors, *C. elegans* represents a powerful model organism to study the fundamental functions of neural circuits (1). The comprehensive mapping of the worm’s connectome has led to many foundational insights into the dynamics by which sensory inputs generate motor outputs (2–4). However, fundamental questions remain unanswered in identifying the dynamics of neural circuits that underlie behavior. We are interested in employing mathematical methods to address the role of multisensory integration in generating chemotactic behavior (5).

Chemotaxis toward a chemical attractant—such as the volatile odorants produced by a food source—is an essential part of the *C. elegans* behavioral palette (6–10). Extensive experimental research has demonstrated that the ability of *C. elegans* to locomote toward a food source depends on a variety of locomotory behaviors, including gradual turning (weathervane) locomotion (7), sharp turns, including pirouettes, and reversals (11). Particularly reversals, i.e. bursts of backward locomotion (12), and omega turns, in which the body of the animal bends so that the head touches the tail (12), are shown to be dependent on the time derivative of the sensory input (11, 13). Efforts to mathematically model nematode chemotaxis have produced insights into how sensory neurons and interneurons encode chemical concentration gradients (7, 11, 14, 15). In the present study, we investigate whether and how a minimal neuronal circuit can integrate sensory information so that its dynamics can produce realistic biomechanical activity, ultimately resulting in successful chemotaxis.

We present a functionally constrained dynamical mathematical model of *C. elegans*, which includes a head circuitry with sensory, inter-, and motor neurons, coupled with a neuromechanical body model with muscles and stretch receptors. Our model worm displays forward locomotion toward a chemical attractant, steered by continuous shallow turns and enriched with strategically timed sharp/omega-like turns that, combined, orient the worm toward the food source. We demonstrate that the emergent ability to perform sharp turns enables chemotaxis in a wide variety of chemical gradients and is facilitated by the integration of chemosensory and proprioceptive cues in the interneuron layer. Our work corroborates experimental findings, and it generates new testable hypotheses about the role of multisensory integration in achieving a dynamic range of behaviors in a compact nervous system such as that of *C. elegans*.

## 2. Results

### 2.1 A neuromechanical model of chemotaxis with emergent sharp turning behavior

The neuromechanical model described herein is based on the one presented in (16) which we have extended to include one sensory neuron (AWA) and two interneurons (RIM and RIA) **(#Figure 1A, see Methods)**, partially following the circuit logic in (17) and in (5), while introducing a different modeling strategy. In the living worm’s connectome, AWA is indirectly connected to RIM through a network of interneurons, including AIY and AIB. The net influence of these pathways on RIM depends on the circuit state and the pattern of sensory input. In our circuit, we have omitted explicit representations of these interneurons while maintaining the condition-dependent dynamics and functional consequence, as changes in AWA activity can either inhibit or disinhibit RIM, depending on the direction of the sensory input change. Thus, the AWA - RIM relationship is interpreted as a reduced functional pathway rather than as a direct anatomical connection. Similarly, in living *C. elegans*, RIM, RIA, SMD, and RMD are embedded in a more extensive network of interneurons and motor neurons. RIM influences the head motor circuit through both synaptic and neuromodulatory pathways, while RIA receives convergent sensory and motor-related input and is reciprocally connected with SMD and RMD. Our reduced model does not represent these intermediate pathways individually. Instead, our model captures their effective dynamical interactions through the coupling between RIM, RIA, SMD, and RMD. The connectivity therefore prioritizes functional and dynamical correspondence with the biological circuit over one-to-one anatomical fidelity. SMD and RMD neurons innervate head and neck muscles and contribute to drive sinusoidal locomotion that begins with head turning and propagates through the body. Downstream body wall muscles are innervated by ventral nerve cord neurons. Across a cycle of locomotion, the worm proceeds forward when dorsal and ventral turns are symmetric. Conversely, changes in direction occur when dorsal and ventral turns are asymmetric.

**Figure 1.**
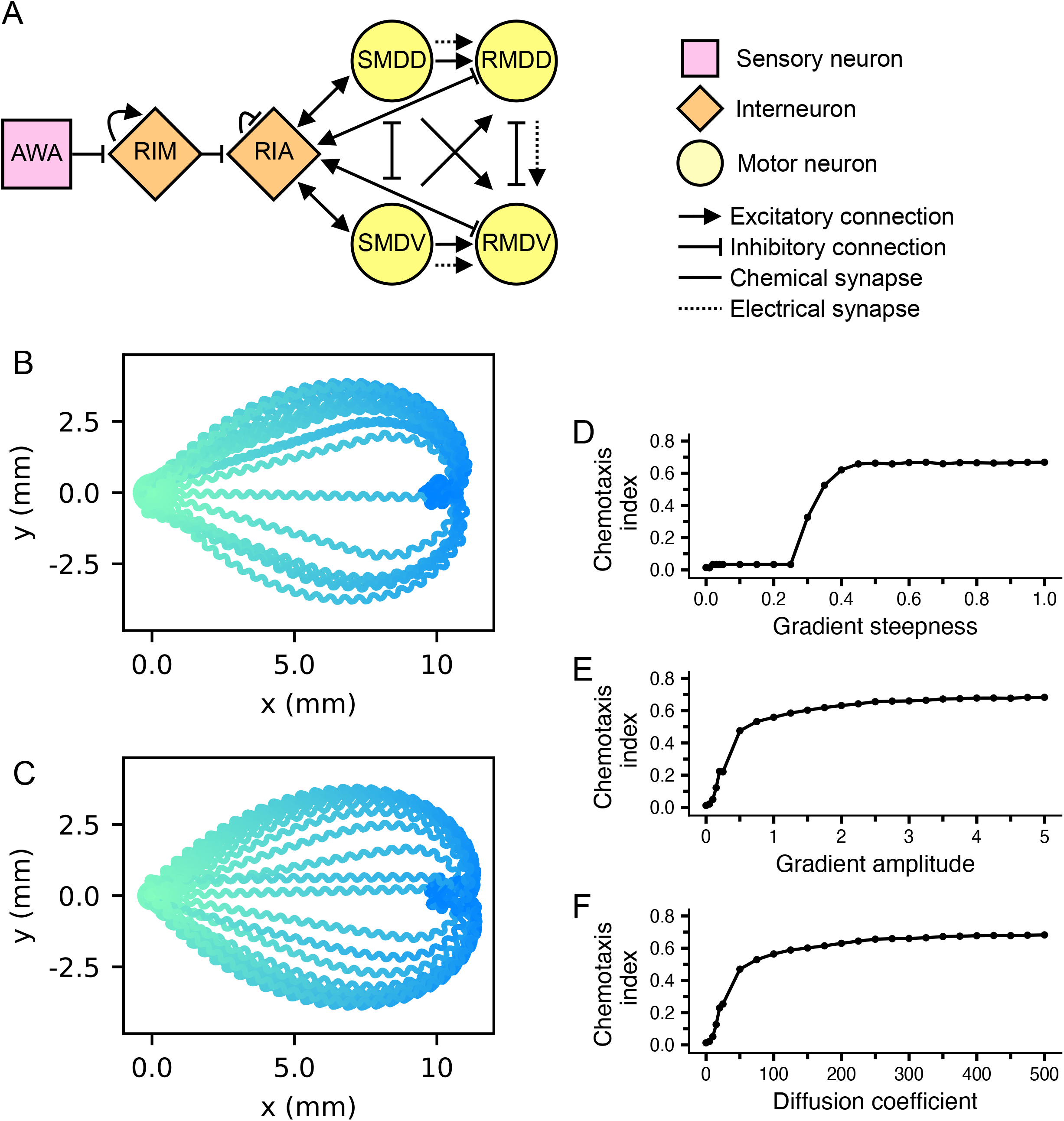
Neuronal circuit model and chemotactic behavior. (A) Head neuronal circuit architecture. (B, C) Behavioral traces of model worm locomotion in a (B) conical and (C) Gaussian attractant concentration gradient field, with varying initial heading angles of [0°, 360°]. Initial distance from food source is 10mm and simulation is run for 100s. Worm path is color- coded according to time step, from green (beginning of simulation) to blue (end of simulation). (D, E, F) Chemotaxis index at varying (D) conical gradient steepness, (E) Gaussian gradient amplitudes, and (F) Gaussian gradient diffusion coefficients. Data is averaged from four simulations with different food positions, placed 10mm from the worm’s initial position with an angle difference of 0°, 90°, 180°, and 270° from the worm’s initial heading angle.

#### 2.1.1 Model parameter optimization

Model parameters were optimized using an evolutionary algorithm with the model worm placed in an attractant concentration field with a point food source and conical gradient with a steepness coefficient of 0.75 (see Methods for details). The optimal parameter set used for analysis, as well as parameter ranges generating optimal solutions for head neuronal parameters, are indicated in **(#Table 1)**. After optimization, the value of the chemotaxis index for model worms in a conical and in a Gaussian attractant concentration gradient field was 0.756 ± 0.006 and 0.702 ± 0.008, respectively. We tested the generalization of the optimized solution on conical gradients of variable steepness and on Gaussian gradients of variable amplitudes and diffusion coefficients **(#Figure 1B–F)**. We found that chemotaxis index, velocity, and turning angle **(#Supplementary Figure 1)** were consistent across a variety of gradients. This generalization of the solution across multiple types of chemical concentration gradients indicated a chemotaxis model suitable for further analysis.

**Table 1.**
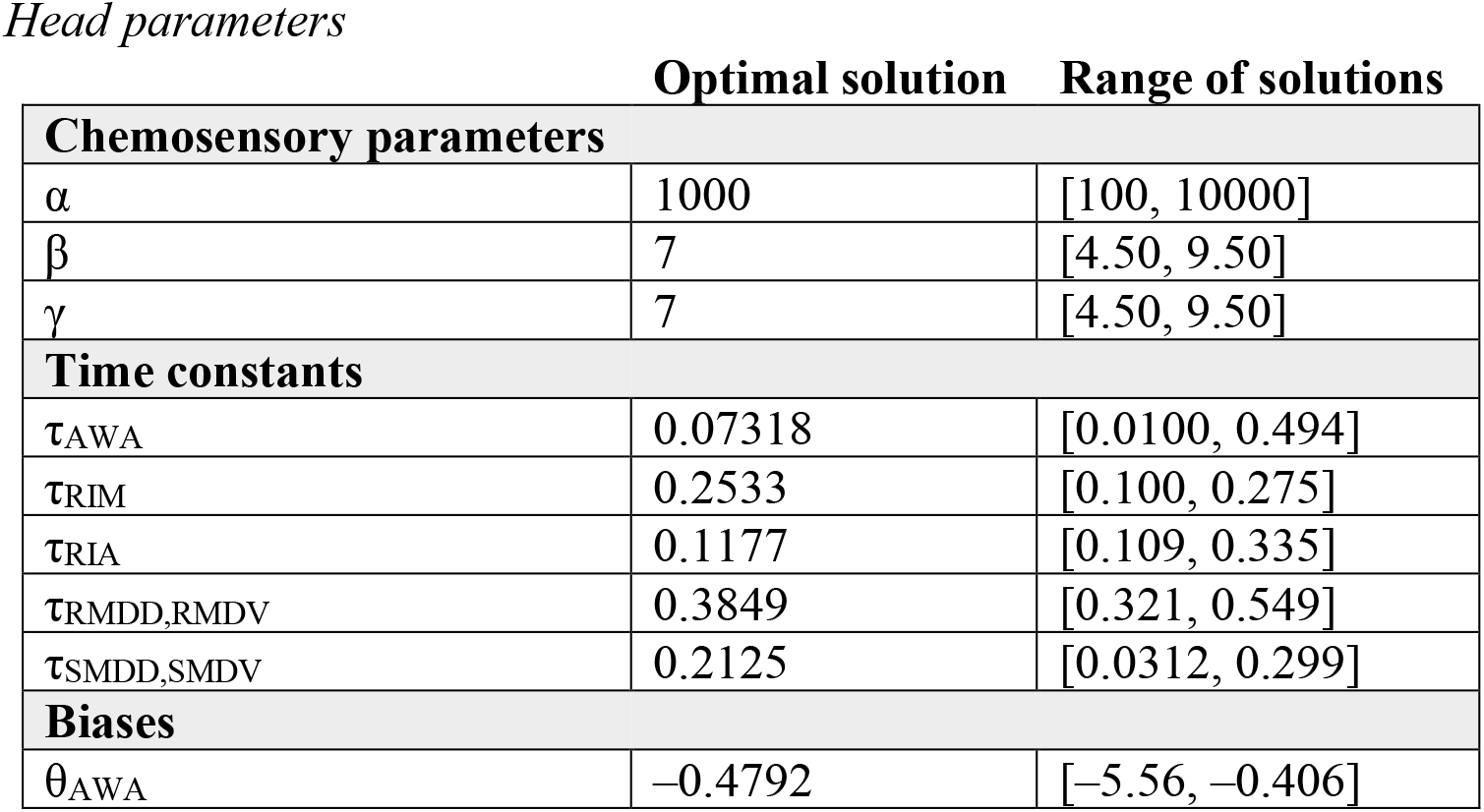
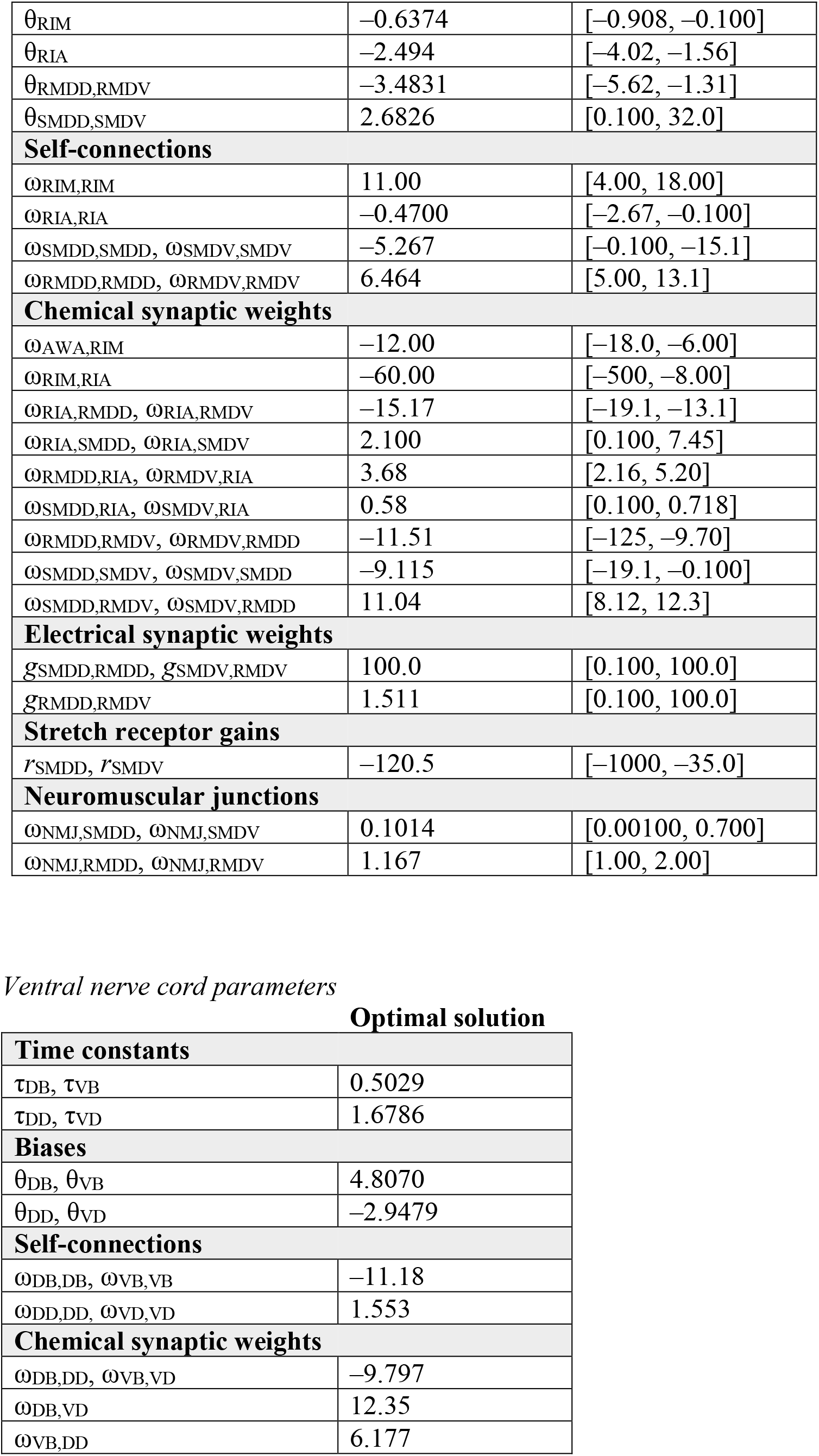
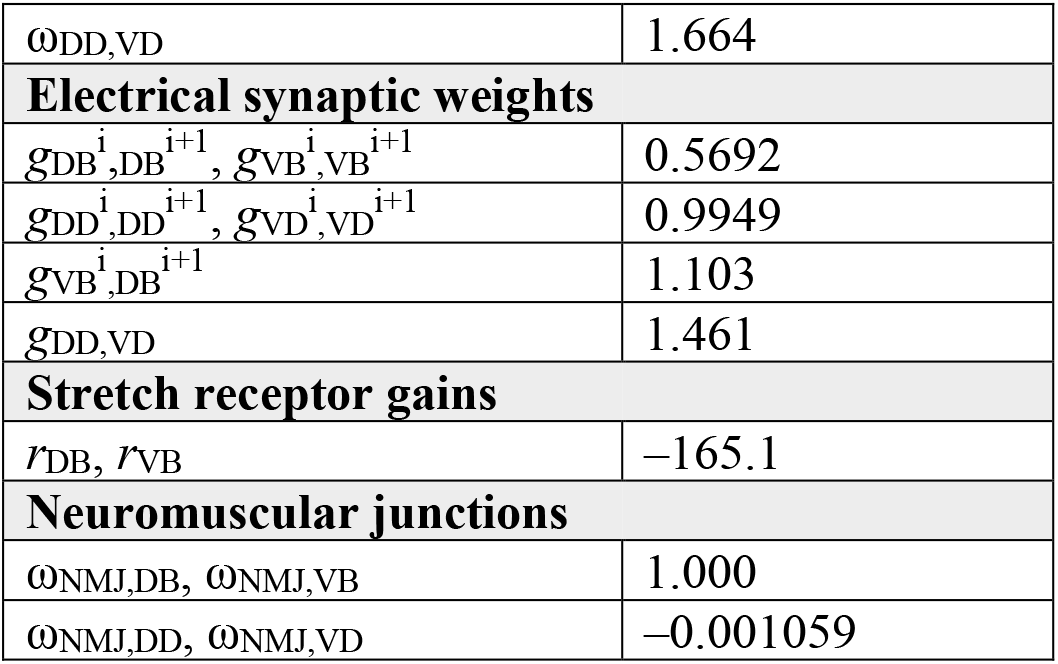
Optimized model parameters.

#### 2.1.2 Sharp turns as an emergent property

The chemotaxis exhibited in this model is accompanied by the worm’s ability to produce both shallow and, strikingly, sharp turns as it navigates toward the food source (**#see frames [0, 1900] of Video 1, [0, 1900] of Video 2, and [0, 2000] of Video 3**). Maximum turning angle is defined in this study as the highest magnitude of a dorsal or ventral turn exhibited by the worm during a simulation (see **Figure 2** for terminology). Before optimization, maximum turning angle was 88.24°. After optimization, maximum turning angle was 272.82°. The ability to produce sharp turns is an emergent property of the model. It occurs during two key model behaviors, both of which ultimately aid chemotaxis. First, sharp turning appears to accommodate rapid initial adjustment to the attractant gradient, regardless of initial heading angle. For instance, when the food source is located 180° from the worm’s initial heading angle, the worm completes a sharp turn at the beginning of the simulation run, which results in rapidly orientation toward the line of steepest ascent **(#see frames [80, 240] of Video 2 and [80, 250] of Video 4**). Second, once the worm reaches the food source, it performs sharp turns that assist with it remaining proximal to the food source for the remainder of the simulation **(#see frames [1900, 2500] of Video 1, [1900, 2500] of Video 2, [1300, 2500] of Video 3, and [2000, 2500] of Video 4)**. Therefore, sharp turns are an integral part of the model’s effective chemotaxis.

**Figure 2.**
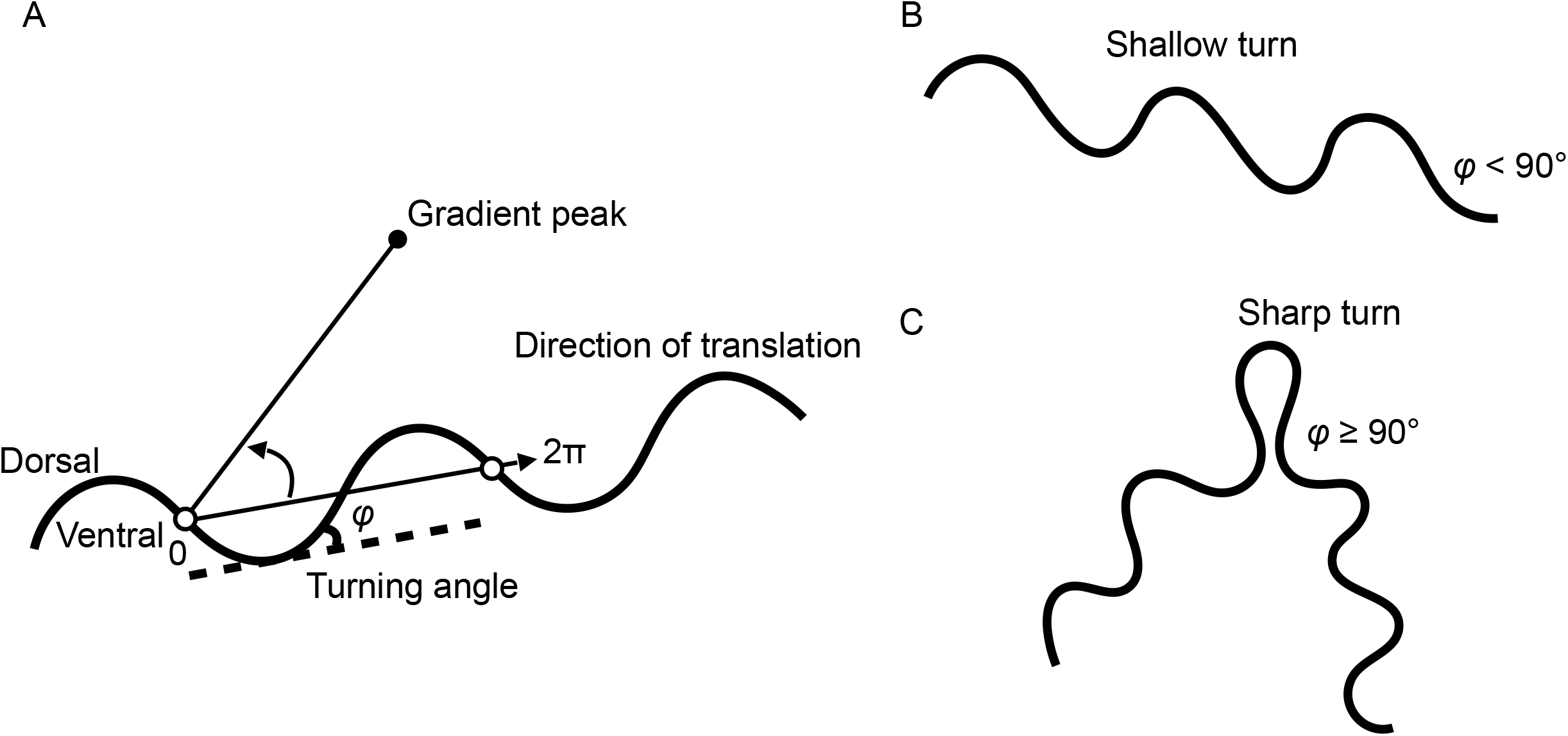
Terminology. (A) Definition of turning angle, one cycle of locomotion (0 to 2π), direction of translation, and bearing. The direction of translation is defined by any two points in the worm’s trajectory separated by a distance of 2π (i.e., the line connecting the current head position with the head position at the same phase of the previous cycle of locomotion). Turning angle (φ) is defined as the angle difference between the direction of translation and the position of the head. Turning bias is defined as the sum of the turning angle across one cycle of locomotion, and bearing is defined as the angle difference between the direction of translation and gradient peak. (B) Definition of a shallow turn (φ < 90°), with example trajectory. (C) Definition of a sharp turn (φ ≥ 90°), with example trajectory of worm completing one sharp turn.

When we examined the body shape of the worm during these sharp turns, we saw that the head touches or almost touches the tail (**#Supplementary Figure 2**), in a configuration similar to the well-known omega turns (18–20).

#### 2.1.3 Sharp turns are triggered by decreases in concentration

We next asked whether sharp and shallow turns are part of a binary turning behavior, dictated by a threshold, or whether sharp and shallow turns smoothly succeed each other, forming a continuous spectrum of varying turning angle magnitudes. To address this, we examined turning bias as a function of bearing **(#see Figure 2 for terminology, and #Figure 3A, B)**. This experiment demonstrated that turning bias has a positive linear relationship with bearing **(#Figure 3B**, r = 0.987). We thereby concluded that the model worm adjusts its turning angle continuously to orient itself in the direction of steepest ascent.

**Figure 3.**
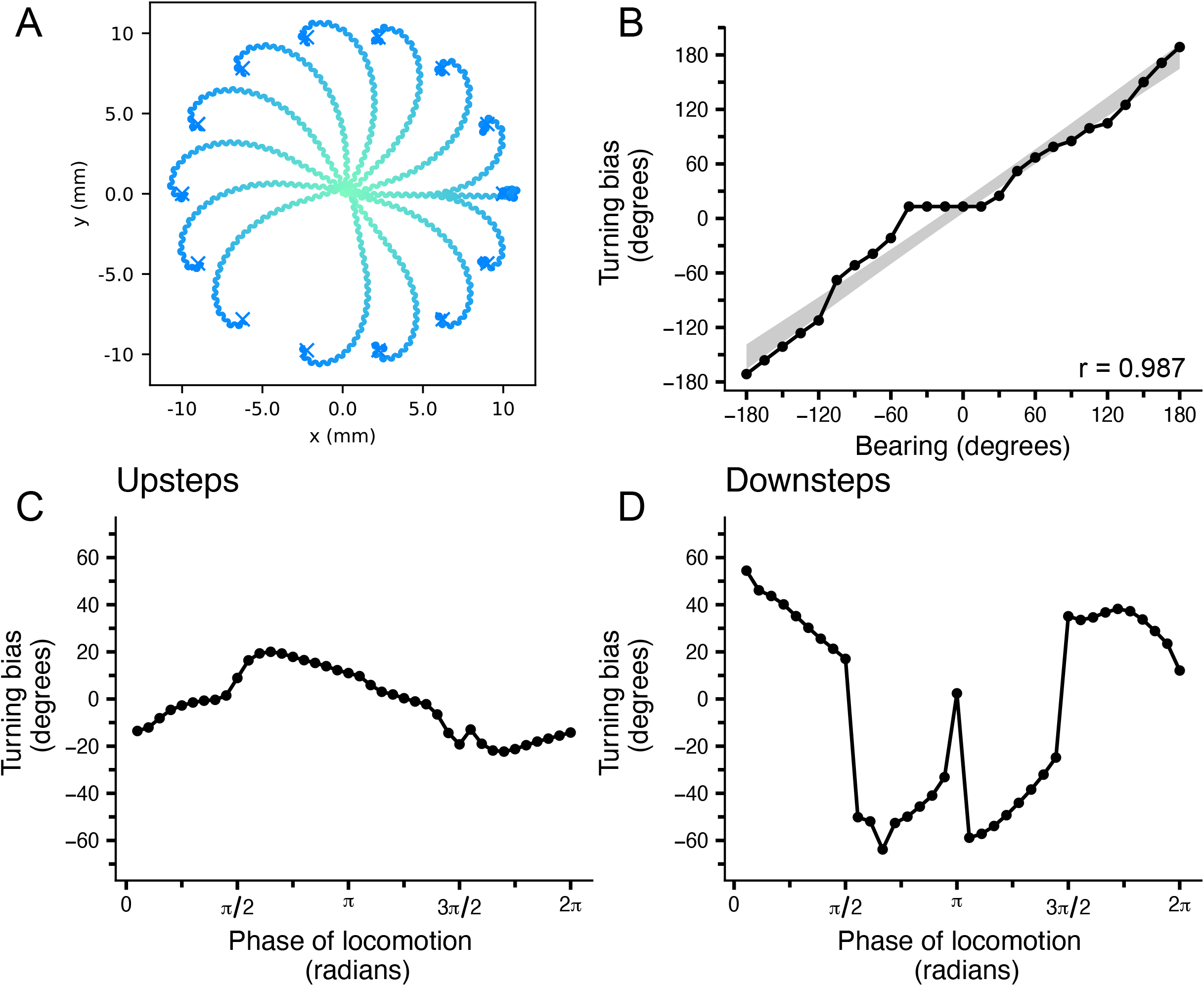
Analysis of chemotaxis. (A) Traces of worms locomoting toward different food source positions placed 10mm from the worm’s initial position of (0.0, 0.0), with time progression indicated as a green–blue gradient. (B) Turning bias as a function of bearing. Turning bias is recorded across one cycle of locomotion following an initial bearing of [–180°, 180°]. Concentration step assay performed during (C) up-steps and (D) down-steps, in which concentration steps either up to +1.0 or down to –1.0 from an initial value of 0.0 at a specific phase of locomotion; turning bias was recorded across the following cycle of locomotion.

Next, we investigated the environmental stimulus that elicits shallow and sharp turning. To this end, we conducted a concentration step assay, in which a concentration step up to +1.0 (up-steps) or down to –1.0 (down-steps) from an initial value of 0.0 is artificially introduced at a specific phase of locomotion, and the resulting turning bias across the cycle of locomotion following the concentration step is recorded.

We found that, in the case of up-steps (**#Figure 3C**), turning bias exhibits a non-monotonic pattern with low magnitude variation around zero mean across a cycle of locomotion. In the case of down-steps (**#Figure 3D**), turning bias exhibits higher magnitude variations across a cycle of locomotion.

This indicates that the model is able to adjust its turning behavior in a phase-sensitive manner. Furthermore, this demonstrates that shallow turns are elicited by increases in concentration, and sharp turns are elicited by decreases in concentration. This demonstrates that the environmental stimulus for sharp turns is the sign of the change in concentration.

#### 2.1.4 Direction of sharp turn depends on phase of locomotion

Though turning bias in both up-steps and down-steps of the concentration step assay shows a dependence on the phase of locomotion (**#Figure 3C, D**), an important distinction emerges in the case of down-steps. Namely, the abrupt difference in turning bias that occurs around π/2 and 3π/2––when the worm is halfway through a dorsal or ventral turn respectively–– indicates that the direction of a sharp turn depends on the phase of locomotion in which the concentration decrease occurs. If a decrease in concentration occurs before a worm has completed half of a turn (i.e., between 0 and π/2, or π and 3π/2), then the sharp turn will take place in the same direction it was bending when the concentration step occurred **(#Figure 3D**, phases of locomotion [0, π/2] and [π, 3π/2]). Conversely, if a decrease in concentration occurs after a worm has completed half of a turn, then the sharp turn will take place in the opposite direction the worm was bending when the concentration step occurred **(#Figure 3D)**. We interpreted this to mean that while sharp turning is triggered by decreases in concentration, the direction of the sharp turn depends on the orientation of the worm when the decrease occurs.

### 2.2 AWA activity is concentration- and history-dependent

We next analyzed AWA neuronal activity to investigate how this neuron encodes changes in concentration to elicit directed turning behavior. In living nematodes, AWA activity has been demonstrated to be concentration- and history-dependent (21). In our model, because the computational instantiation of AWA output is dependent on both fast and slow sensory inputs, the temporal differentiation of AWA input could underpin the model’s behavioral response to changes in concentration.

To explore this, we considered specific concentration gradients and recorded sensory inputs as well as AWA output **(#Figure 4)**. We chose to deliver two types of constant concentrations—at either +1.0 or 0.0 **(#Figure 4A, B)**—as well as two types of variable concentration—either linearly increasing from 0.0 to +1.0 or linearly decreasing from +1.0 to 0.0 **(#Figure 4C, D)**—and then recorded sensory inputs to AWA and its output across 1 second in order to determine how AWA differentially encodes changes in concentration and static concentration of the same magnitude.

**Figure 4.**
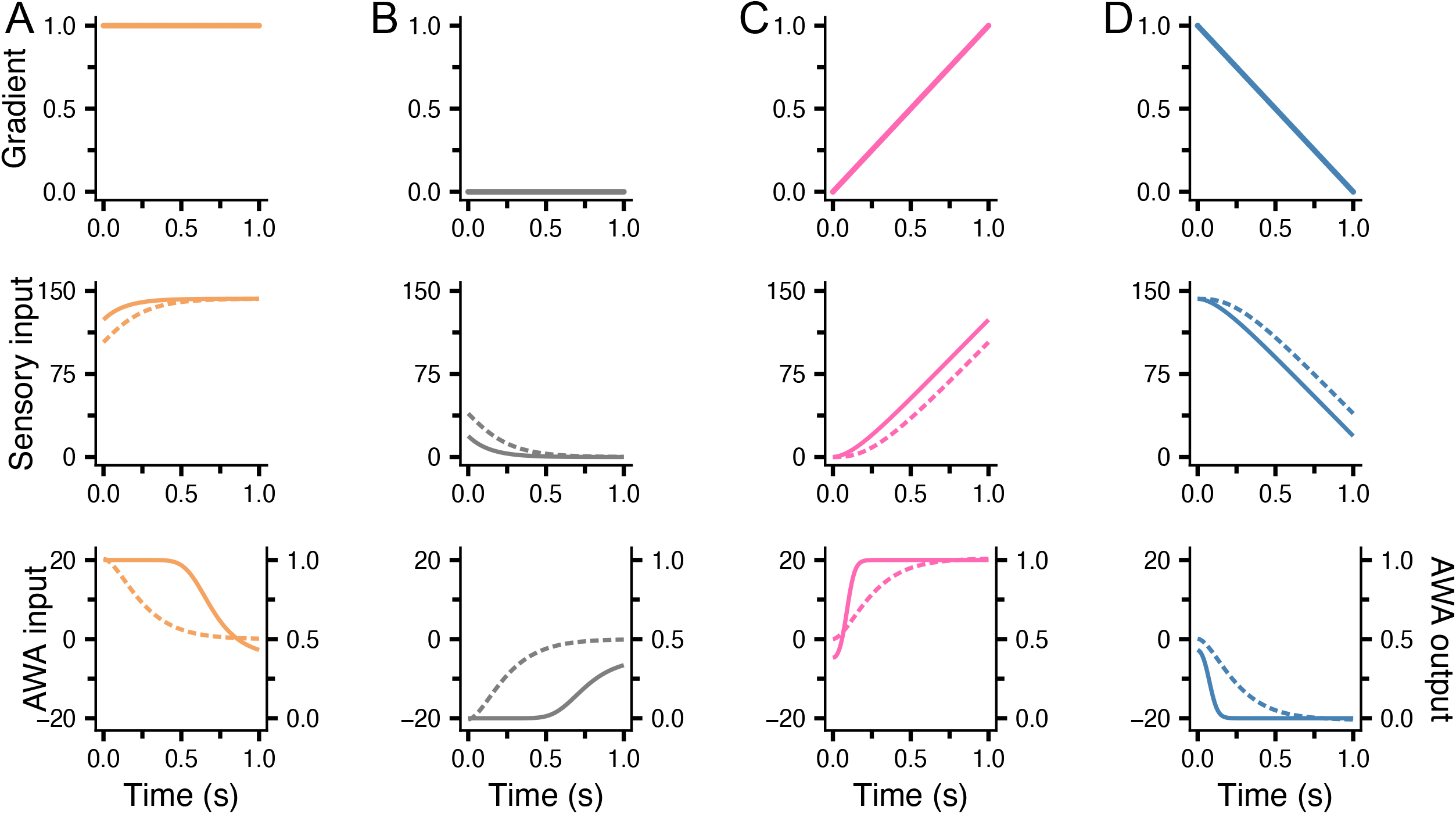
Analysis of AWA activity. Artificial induction of either a (A, B) constant, (C) linearly increasing, or (D) linearly decreasing concentration across one second. Concentration varies in the range [0.0, 1.0] or is static at {0.0, 1.0}. Top row depicts value of concentration. Middle row depicts fast (solid) and slow (dashed) sensory inputs. Bottom row depicts AWA input (dashed, left axis) and output (solid, right axis).

In the case of constant concentration, fast and slow sensory inputs converge **(#Figure 4A, B)**. This results in AWA output approaching a value of 0.382 within 3.48 ± 0.0423 seconds both when concentration is constantly high **(#Figure 4A)** and when it is constantly low **(#Figure 4B)**. Conversely, when concentration changes linearly, fast and slow sensory inputs diverge **(#Figure 4C, D)**. This results in AWA output either ramping to +1.0 **(#Figure 4C)** or diminishing to 0.0 **(#Figure 4D)** within 35.5 ± 8.50 milliseconds in response to increases and decreases in concentration, respectively. This means that AWA rapidly maximizes its output in response to increases in concentration, rapidly minimizes its output in response to decreases in concentration, and slowly plateaus its activity in response to constant concentration. Given that AWA is the only chemosensory neuron in the model, these dynamics, in sum, form the initial encoding that steers chemotaxis.

### 2.3 RIM modulates RIA activity to promote sharp turns in response to decreases in concentration

We next analyzed the dynamics of RIM and RIA activity in the context of shallow and sharp turning to determine how these neurons encode AWA activity to elicit chemotaxis.

In a living worm, RIA activity tends to oscillate in time with head swings, which promotes gradual readjustment toward the line of steepest ascent in the concentration gradient field, as is observed in klinotaxis (22, 23). At the same time, RIM serves as a state switch to elicit reversal behavior in response to decreases in chemical concentration (24). This activity allows rapid re- orientation toward the line of steepest ascent, as is commonly observed in klinokinesis and pirouette behavior (11). Though our model captures forward locomotion rather than reversal- including pirouette behavior in chemotaxis, we sought to analyze RIM activity in the context of omega/sharp turns as sharp turning is recapitulated in our model.

We hypothesized that RIM signaling promotes sharp-turn chemotaxis in response to decreases in concentration, while RIA signaling elicits shallow-turn chemotaxis in response to increases in concentration. To test this, we plotted turning angle as well as AWA, RIM, and RIA activity in the context of shallow-turn and sharp-turn chemotaxis **(#Figure 5)**. In the context of shallow-turning behavior **(#Figure 5A)**, brief decreases in concentration cause rapid and brief decreases in AWA activity within 51.85 ± 3.64 milliseconds. This results in RIM either producing no activity, or spiking briefly at a low magnitude when AWA is sufficiently low (see Figure 5A at 6.7 seconds and 13.4 seconds), which inhibits RIA briefly. This, in turn, causes a brief disruption in RIA oscillations from 0.606 ± 0.00218 Hz (in regular straight-line motion) to 0.507 ± 0.0440 Hz (computed across the interval of [5, 15] seconds in Figure 5A), an insignificant change, t(6) = 2.25, p = 0.109), thereby allowing a gradual, shallow turn toward the food source.

**Figure 5.**
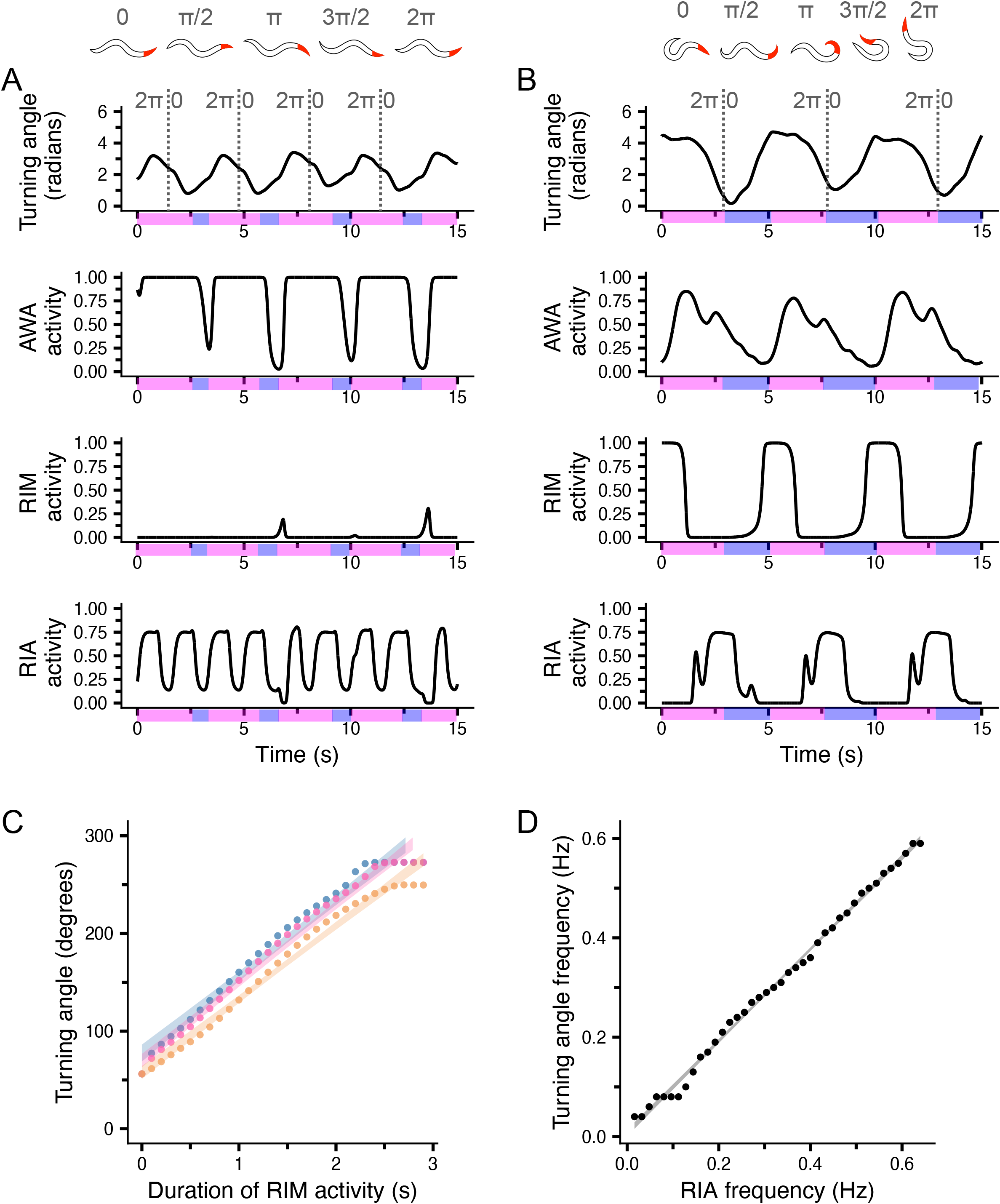
AWA–RIM–RIA circuit dynamics in shallow and sharp turning. Turning angle and neuronal activity from AWA, RIM, and RIA in (A) shallow- and (B) sharp-turn chemotaxis. Schematics of worm body positions during shallow and sharp turns across a cycle of locomotion are given above graphs with red indicating the head position of the worm. Dotted grey lines indicate the time points at which a cycle of locomotion begins. Magenta shading indicates time steps at which concentration is increasing (Δc > 0). Blue shading indicates time steps at which concentration is decreasing (Δc < 0). (C) Turning angle as a function of RIM activity duration. Activity is held at a constant value of either 1.0 (blue), 0.5 (red), or 0.1 (yellow) for ≤ 3s, and the magnitude of the resulting turn in one given direction is recorded. (D) Turning angle frequency as a function of RIA activity frequency. RIA frequency varies from [0 Hz, 0.6 Hz] for 50s., and resulting turning angle frequency is recorded across the entire simulation (for reference, RIA frequency is 0.6 Hz when the worm is moving in a straight line). For both (C) and (D), linear regression was performed and the confidence intervals of the lines of best fit are included in the graphs. Correlation coefficients are as follows: (C) when I_RIM_ = 1.0 (blue), r = 0.991; I_RIM_ = 0.5 (red), r = 0.986; I_RIM_ = 0.1, r = 0.991. (D) r = 0.998.

Conversely, in the case of sharp turning **(#Figure 5B)**, a prolonged decrease in concentration causes AWA activity to diminish from its maximum value to its minimum value within 3.385 ± 0.186 seconds. The area under the curve (AUC) of AWA activity across a 30 second interval decreases from 27.16 ± 0.67 to 12.39 ± 0.017 **(#Supplementary Figure 3)**. This causes RIM activity to spike strongly and remain activated for longer periods of time. The AUC of RIM activity across a 30 second interval increases from 1.10 ± 0.40 to 9.79 ± 0.12 **(#Supplementary Figure 3)**. The resulting increased inhibition of RIA lowers the frequency of RIA oscillations to 0.371 ± 0.0315 Hz, significantly lower than RIA oscillation frequency in shallow turning (t(6) = 7.43, p = 0.0049) which ultimately produces a sharp turn.

In order to determine the role of RIM and RIA activity in sharp turning behavior, we next analyzed turning angle as a function of RIM activity duration **(#Figure 5C)** and RIA activity frequency **(#Figure 5D)**. To do so, we induced a RIM current of variable strengths and duration at the beginning of a head-led turn and recorded the magnitude of the following turning angle.

We found that turning angle magnitude increases linearly as a function of RIM activity duration **(#Figure 5C**, r = 0.989). We next induced a sinusoidal RIA current of variable frequencies and measured the resulting frequencies of head turning. We found that turning angle frequency increases linearly as a function of RIA activity frequency (**#Figure 5D**, r = 0.989). A lower frequency turning angle allows a turn to occur across a longer period of time, thus allowing a sharp turn as the magnitude of turning angle increases over time. These data suggest that the concerted action of RIM activity duration and RIA activity frequency modulates turning angle magnitude and frequency to elicit both shallow and sharp turning in chemotaxis.

### 2.4 RIA modulates motor neuron activity to produce sharp turns

We next investigated how the modulation of RIA oscillations coordinates downstream neuronal activity and changes in kinematics during chemotaxis. To do so, we measured the activity of RIA, RMD and SMD motor neurons, head stretch receptors, and head muscles in the context of shallow and sharp turns **(#Figure 6)**.

**Figure 6.**
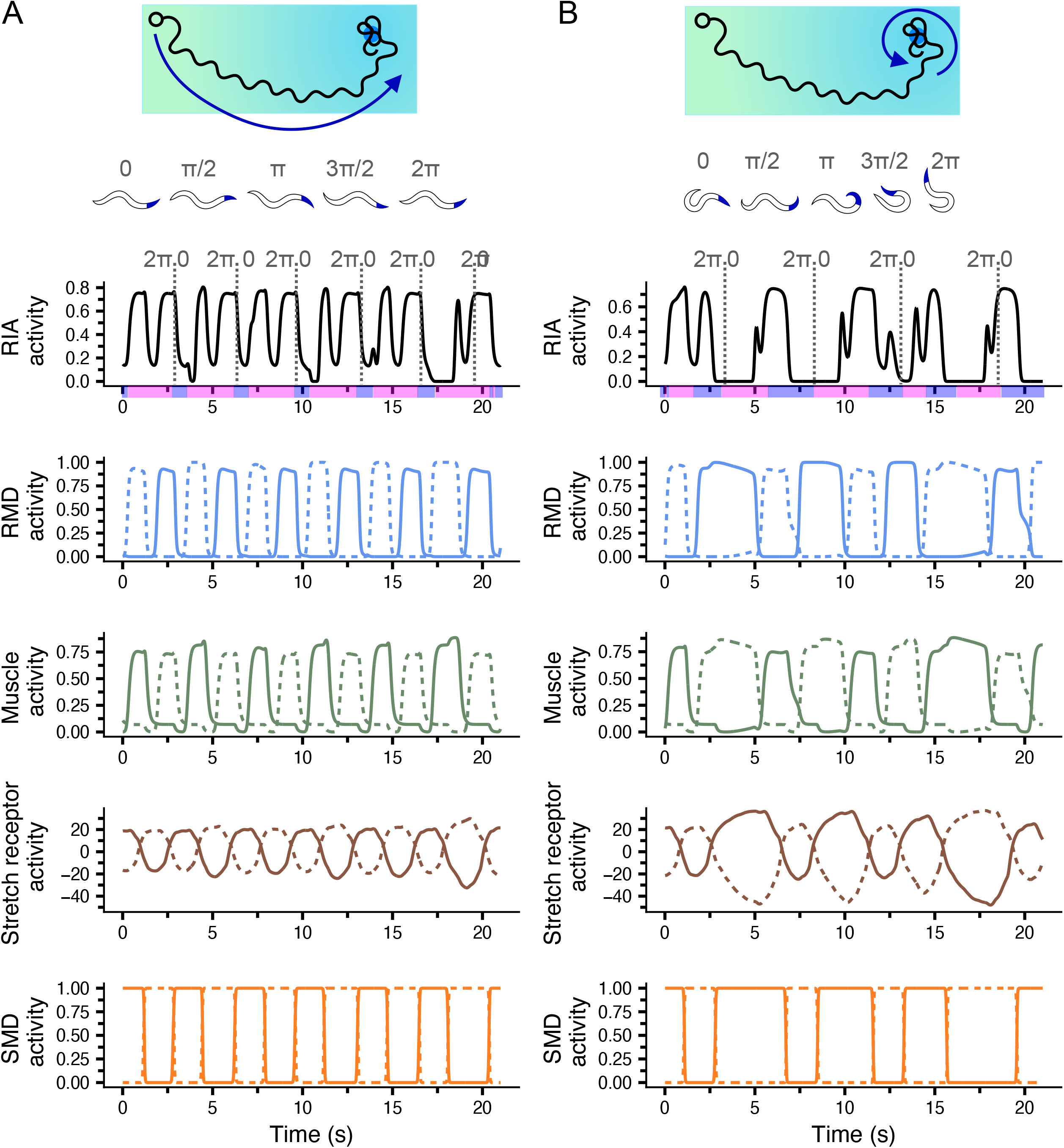
Stretch receptors’ dynamics shape sharp turning behavior. RIA, stretch receptor, motor neuron, and muscle activity in (A) shallow turns and (B) sharp turns. Both graphs are generated from a 100 second simulation; (A) shows outputs over a 20 second interval early in the simulation, when the worm is still far from the food source and performs shallow turns. (B) shows outputs over a 20 second interval late in the simulation, when the worm is close to the food source and performs sharp turns. Top: blue arrow indicates the trajectory of the worm analyzed in each panel. Dotted grey lines indicate where a cycle of locomotion begins, and representative behavioral traces from the worm (head highlighted in blue) across a cycle of locomotion are given. For RIA neuronal activity, magenta shading indicates the time steps at which concentration is increasing (Δc > 0) and blue shading indicates the time steps at which concentration is decreasing (Δc < 0). Dorsal stretch receptors, motor neurons, and muscles are indicated with a solid line, while their ventral counterparts are indicated with a dashed line.

In the context of shallow turns **(#Figure 6A)**, dorsal and ventral head stretch receptor activity is approximately antiphase. SMD dorsal and ventral neurons receive input from ipsilateral head stretch receptors and provide feedback to RIA. SMD and RMD together drive muscle activation, which synchronizes mechanical oscillation with RIA activity. This suggests that stretch receptor and chemosensory oscillations coordinate the kinematics necessary for gradual, shallow turning toward the food source.

Conversely, sharp turns **(#Figure 6B)** are represented in stretch receptors by increases in magnitude (from a maximum amplitude at oscillation peaks of 20.76 ± 0.55 to 35.33 ± 0.85, t(14) = 14.42, p<0.001) and by a decrease in oscillation frequency (from 0.330 ± 0.0152 Hz to 0.165 ± 0.0111 Hz, t(14) = 8.77, p<0.001). These changes are reflected in SMD activity, and, in turn, RMD activity. SMD and RMD input to body muscles promote prolonged asymmetrical muscle activation, which ultimately produces a sharp turn in either the dorsal direction (as seen starting at 2.8 s and 7.1 s of Figure 6B, when RMDD and SMDD activity is prolonged) or the ventral direction (as seen at 15.6 s of Figure 6B, when RMDV and SMDV activity is prolonged).

### 2.5 Muscles and stretch receptors propagate shallow and sharp turns in chemotaxis

#### 2.5.1 Role of muscles in sharp turning and chemotaxis

To explore muscle activity in both shallow- and sharp-turn chemotaxis, we generated kymographs for both cases **(#Supplementary Figure 4)**. These graphs indicate that, throughout chemotaxis, locomotory undulations originate in the head and are propagated posteriorly through the body. Muscles throughout the body support sharp turning by referring oscillatory activity generated by head neuronal circuitry to posterior body segments. When a lower frequency oscillation is propagated through the body, a sharp turn occurs.

#### 2.5.2 Role of proprioceptive information in sharp turning and chemotaxis

To test the role of proprioceptive information in sharp turning behavior, we manipulated stretch receptor activity across a concentration step down **(as in #Supplementary Figure 1)**.

When stretch receptor activity was allowed to respond to upstream activity **(#Supplementary Figure 5A)**, the silencing of RIA activity following a concentration step caused RMDD activity to be prolonged, which is reflected in stretch receptors by a prolonging of their activity and increase in magnitude. This then activates SMDD, which activates RMDV to end the current dorsal sharp turn and begin a ventral sharp turn.

In this assay, stretch receptor activity is manipulated to oscillate in a sinusoidal pattern equal in frequency to shallow turning in the model **(#Supplementary Figure 5B)** to disrupt proprioceptive feedback. Following the concentration step at the beginning of a dorsal turn, RMDD is activated, which then activates dorsal muscles. This produces a dorsal sharp turn.

However, SMD neurons continue to oscillate in time with stretch receptors. This leads to a failure in contralateral RMD activation following a sharp turn, and locomotion stalls as both RMDD and dorsal muscles remain persistently active. When the concentration step ends, locomotion proceeds normally after a delay. In our model, stretch receptor activity is necessary to engage SMD, which excited the contralateral RMD, thereby promoting rapid re-adjustment between dorsal and ventral turns.

### 2.6 Effect of *in silico* synaptic ablation on sharp turning and chemotaxis

To determine the contributions of specific synaptic connections within the head neuronal circuitry to shallow- and sharp-turn chemotaxis, we next explored changes in the chemotaxis index and maximum turning angle of the model worm when targeted chemical synapses were ablated **(#Figure 7)**.

**Figure 7.**
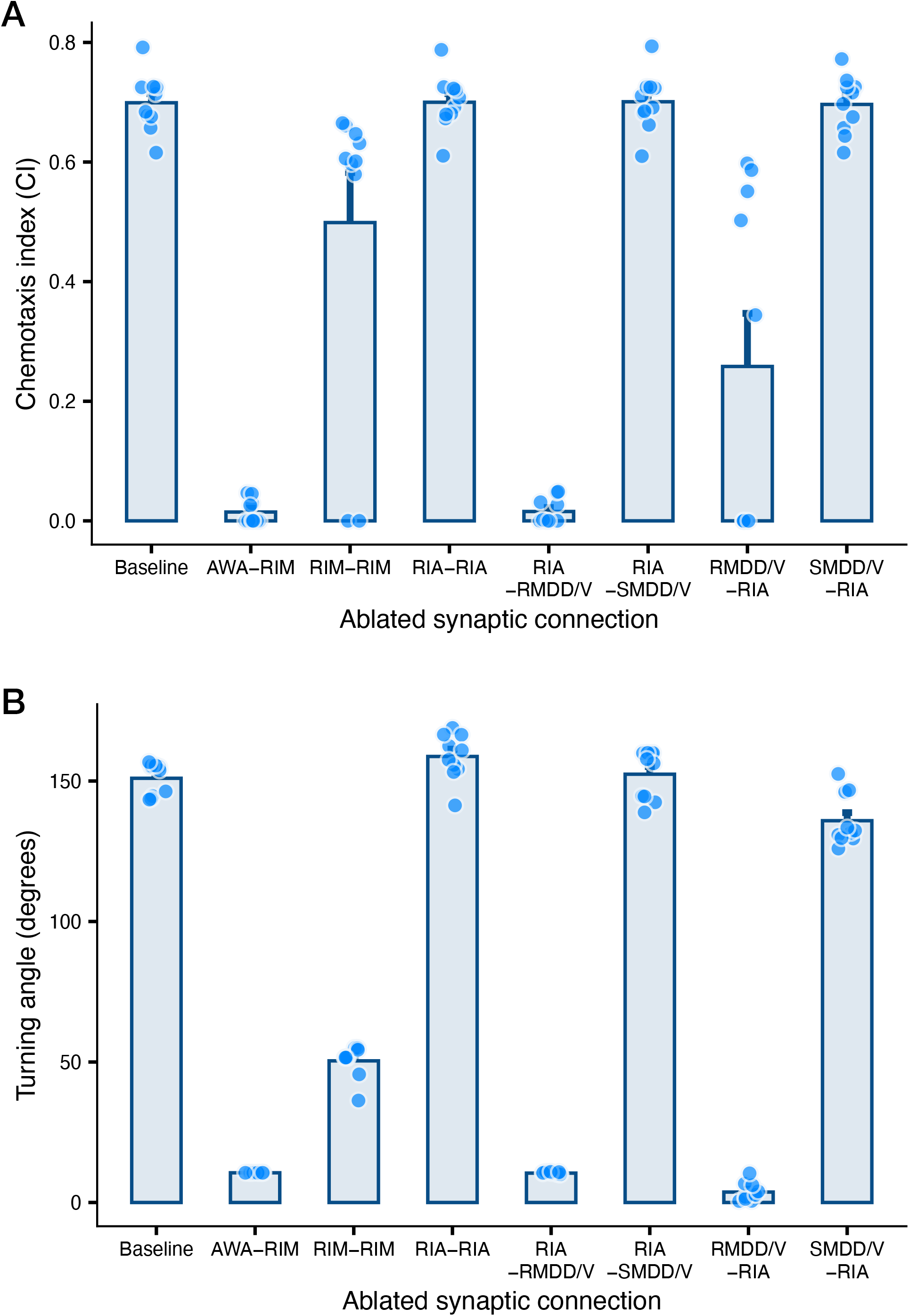
Effect of simulated synaptic ablation in chemotaxis and sharp turning behavior. For each manipulation, chemotaxis index or turning angle is recorded across ten simulations in which the model worm has the same initial heading angle and variable food locations. Individual points represent the values from each individual simulation. Bar plots show the mean value of each condition with SEM. (A) Chemotaxis index with all chemical synapses present (baseline) and with specific chemical synaptic ablations, including ω_AWA–RIM_; ω_RIM–RIM_; ω_RIA–RIA_; ω_RIA– RMDD_, ω_RIA–RMDV_; ω_RIA–SMDD_, ω_RIA–SMDV_; ω_RMDD–RIA_, ω_RMDV–RIA_; and ω_SMDD–RIA_, ω_SMDV–RIA_. (B) Turning angle in the worm at its most proximal distance to the food source with specific chemical synaptic ablations.

Both the baseline fitness and the fitness when AWA–RIM is ablated are given as references. Overall, one-way ANOVA revealed a significant effect on chemotaxis index, F(7, 63) = 47.96, p < 0.0001. Intriguingly, silencing RIA–RMD (0.0156 ± 0.00658) or RMD–RIA (0.258 ± 0.0888) has a larger negative impact on chemotactic fitness than silencing RIA–SMD (0.701 ± 0.0152) or SMD–RIA (0.696 ± 0.0152) **(#Figure 7A)**.

These ablations also revealed a significant effect on maximum turning angle, F(7, 63) = 1371, p < 0.0001. Silencing SMD–RIA (136 ± 2.87), RIM–RIM (50.4 ± 1.78), AWA–RIM (10.6 ± 0.001), RIA–RMD (10.5 ± 0.110), and RMD–RIA (3.72 ± 1.00) significantly lowered maximum turning angle compared to baseline (151 ± 1.79), while silencing RIA–RIA (159 ± 2.59) or RIA–SMD (152 ± 2.74) did not significantly affect performance **(#Figure 7B)**. The importance of the RIM self-connection in turning angle magnitude may indicate the importance of this neuron’s spiking activity in sharp turning. The increased sensitivity of the model to ablation connection between RIA and RMD compared to RIA and SMD may be due to the network architecture (see Discussion).

## 3. Discussion

We have developed and characterized an integrated neuromechanical model for *C. elegans* chemotaxis (**#Figure 1**) that exhibits sharp turning behavior as an emergent property (**#Figure 3, Supplementary Figure 2,** and **Videos 1- 4)**. The model is based and builds on prior well-established mathematical modeling (5, 16, 17, 25–30) and experimental work (15, 18, 31). In parallel, our model prioritizes conceptual representations of neuronal connections (e.g., AWA to RIM, RIM to RIA) that account for context-dependent abstraction of real circuitry dynamics, instead of connectome-based individual neuronal connections. This modeling strategy, combined with a tailored optimization approach, allows the model worm to respond dynamically and realistically to a chemical attractant gradient. This includes the performance of sharp turns that resemble a living worm’s omega turns (**#Figure 2, Supplementary Figure 2, and Videos 1- 4**), without the need for an explicit or hardcoded mathematical component (32, 33). Together, sharp and shallow turns enable the model worm to stay on track and correct its path toward the food source, as is the case with living nematodes (6, 7, 9, 11, 18, 21, 34, 35).

It is well established that, during *C. elegans* chemotaxis, the largest change in direction is generated via an omega turn, during which the animal’s body shape resembles the Greek letter Ω (12, 18). In our model, sharp turns (φ≥90°, **#Figure 2C**) occur at the beginning or at the end of the worm’s path, at instances when the worm moves down the chemical gradient (**Videos 1-4**). During these sharp turns the body of the worm is coiled so that the head almost touches or touches the tail or slightly crosses the posterior body part (**Supplementary Figure 2**). To our knowledge, this is the first time that omega-like turns emerge from an embodied chemotaxis navigation model.

The activity of AWA, the only chemosensory neuron in the model (#**Figure 1**), is known from the literature to be both concentration- and history-dependent (21). AWA’s declining response to constant, high concentration of an odorant is referred to as “desensitization” (21, 36). In our model, AWA neuron responds to constant concentration levels, exhibiting a desensitization-like behavior. It also responds to increasing concentration, showing a plateau behavior, in different timescales **(#Figure 4)**. Action potential traces of AWA have shown that AWA can capture small stimulus up-steps by firing immediate spike bursts (37). The plateauing behavior in the model AWA is observed during a continuous and not a step-wise gradient increase (37) and could be attributed to the absence of detailed modeling of the neuron’s ion channels.

We demonstrate that sharp turning is triggered by AWA responses to changes in the concentration of the attractant, in alignment with experimental findings (38, 39). Moreover, sharp turning is locomotion phase-sensitive and is steered by RIM elevated activity and by altered RIA oscillatory activity (**#Figure 5**). A very recent study on the dynamics of olfaction-driven navigation (34) reports findings that are consistent with a pivotal reorientation-promoting role for RIM and provides experimental support for precisely the idea that a sensory gradient can influence *C. elegans* reorientation through the RIM/head-steering circuit. In addition, it has been shown that locomotory response to a sensory stimulus via turning depends on where the worm is in the locomotion cycle, and that Ω-turns are coupled to the dynamics of the existing body-wave oscillator (19). Furthermore, in the model, both shallow and sharp turning are facilitated by changes in head motor neurons (SMDs, RMDs) asymmetric oscillations and supported by asymmetric stretch receptor and muscle activity (**#Figure 6**), as the oscillatory wave propagates through the body (**#Supplementary Figure 3**). It has been suggested that the core mechanism of regulating klinotaxis is the phase-dependent response to sensory stimuli of neurons that steer neck movement (40), and such a role has been proposed for SMDs, based on experiments (41, 42). Recent experimental work further implicates SAA neuron as an intermediate component of the RIM–RMD–SMD turning pathway (34); in the present reduced model, this component is subsumed into the effective coupling between RIM and the downstream head-motor circuit.

In the model, the direction of sharp turns is the result of circuit architecture **(#Figure 6**). In the model, decreases in attractant concentration suppress RIA activity, and the resulting change in RIA dynamics interacts with the phase of the ongoing locomotory oscillation to determine the direction and magnitude of the ensuing turn. A pattern emerges to induce asymmetrical oscillations in motor neuron, stretch receptor, and muscle activity and therefore facilitate sharp turns. First, concentration decreases, which results in RIA activity being silenced.

This prolongs either dorsal or ventral RMD activity, depending on the phase of locomotion in which the concentration step occurs. This RMD activity will engage its respective muscle, which activates stretch receptors. The stretch receptors will then signal the state of the worm’s body to SMD neurons, which causes the SMD neuron ipsilateral to the current turning direction activity to prolong. SMD then increases activity in the contralateral RMD neuron, which allows a sharp turn in one direction to complete and begin the sharp turn in the opposite direction (**#Supplementary Figure 5**).

The finding that silencing the excitatory RIM self-connection has a larger negative impact on chemotactic index and maximum turning angle than silencing the inhibitory RIA self-connection **(#Figure 7B)** could further support the interpretation that positive feedback through RIM amplifies transient changes in RIM activity and promotes transitions between locomotory states associated with sharp reorientation, while RIA activity consistently oscillates with head swings and coordinates more gradual oscillatory changes to promote chemotaxis. Note that the connectivity in the model aims to conceptually capture context-dependent neuronal dynamics and does not stem from explicit connectome-derived architecture. Therefore, the role played by RIM or RIA in our model could be sought in other neurons or collections of such in living worms. The predictions herein for each neuronal component generate plausible hypotheses, aspiring to guide experiments designed to test them.

The ablation experiment (**Figure 7**) revealed variable sensitivity of the model’s chemotaxis and sharp turning behavior to different synaptic perturbations. The difference in sensitivity between manipulating RIA-RMD versus RIA-SMD connections **(#Figure 7)** stems from the neuronal circuit architecture **(#Figure 6**). First, because SMD receives the majority of its total input from stretch receptors **(#Supplementary Figure 6F, 6G)**, RIA will have a disproportionately large effect on RMD activity compared to SMD activity. Second, in our model, RMD neurons receive input from SMD, whereas the reciprocal SMD-RMD connection is not explicitly represented **(#Supplementary Figure 6D, 6E)**. Because of this, silencing the SMD–RIA feedback connection does not functionally ablate all SMD feedback to RIA. This may explain why silencing SMD–RIA and RMD–RIA together has a larger effect on chemotaxis index than silencing RMD–RIA alone (results not shown).

The analysis described herein is based on a deterministic model (43, 44). Nevertheless, we explored the impact of Gaussian noise in individual neurons on the model worms’ chemotaxis index (**#Supplementary Figure 6**), and we found the tolerance of the model to be satisfactory. Because of stochasticity in living systems, further investigating the potential role of noise in a model’s chemotactic and turning behavior constitutes an area of interest.

While we focus on a model parameter set determined by an evolutionary algorithm, we found that the ensemble of models with parameters in the ranges outlined in **(#Table 1)** and in **(#Supplementary Figure 8)** present a qualitative unity. Overall velocity, chemotaxis efficiency, turning frequency, as well as AWA and RIM responses to changes in concentration and the frequency of RIA and motor neuronal correlation to overall turning frequency remain a persistent feature within the ensemble.

The limitations of this model highlight potential areas of future research. First, our model does not capture reversals, since it does not include the respective circuitry (12, 45, 46). RIM has been implicated in reversal initiation and pirouette behavior during chemotaxis (11, 24).

However, because this model is built on a mathematical framework of forward locomotion, we did not attempt to produce reversal behavior here. Second, the inclusion of only a single chemosensory neuron (AWA) was chosen for parsimony but precludes analysis of the multiple chemosensory neurons (AWC, ASE, etc.) present in living worms, and of their synergistic action. Furthermore, AWA in this study reproduces some characteristics of this neuron biologically, but not all-or-none action potentials (37). Overall, while the parsimony of our model aids interpretation and suggests minimum requirements for complex maneuvers, i.e., omega turns, by emphasizing the dynamical aspect, it limits generalization of the findings in the context of other odorants.

To conclude, we present a mathematical framework based on an integrated model of *C. elegans* head chemosensory circuit and body neuromechanical circuit, which successfully recapitulates major chemotaxis features, including sharp/omega-like turns. Rather than reconstructing part of the connectome, the model captures the functional architecture and dynamical interactions of a minimal sensorimotor circuit, in which changes in sensory input alter the dynamical state of RIM, RIA, RMD, and SMD. We show that biologically constrained circuit architecture can generate sharp/omega-like turns as an emergent dynamical consequence of sensory input changes and circuit dynamics, without explicitly programming such elaborate locomotory maneuvers. Our work generates new testable hypotheses about the role of neuronal dynamics and paves the way for the exploration of more complex behaviors (decision making, conflicting cues, multisensory integration) using the suggested modeling approach in conjunction with experiments.

## 4. Methods

The main goal of the proposed framework is to provide a simplified mathematical model for the concerted neuronal and muscle activity that drives *C. elegans* chemotaxis toward a chemical attractant. The model expands upon previously published *C. elegans* models of forward locomotion and chemotaxis (16, 17). To elucidate the computational role of chemosensory input and proprioceptive feedback in driving chemotaxis, the worm is represented by a neuronal circuity consisting of a chemosensory neuron, interneurons, and motor neurons, along with dorsal and ventral body wall muscles and stretch receptors. This model is placed within a two-dimensional chemical attractant concentration gradient to investigate the computational basis of multisensory integration in navigation.

### 4.1 Neuronal circuit architecture

Adapting from a model of *C. elegans* locomotion (16), we consider a neuronal circuit with five main components: (i) the chemosensory neuron AWA, which drives chemotaxis by receiving input from the external environment through chemoreceptor activation, (ii) the interneurons RIM and RIA, the latter of which forms synaptic connections with downstream motor neurons; (iii) the motor neurons RMDD, RMDV, SMDD, and SMDV, which modulate the ventral (V) and dorsal (D) activation of head and neck muscles for forward locomotion, and (iv) a repeating ventral nerve cord (VNC) circuit consisting of B- and D- class motor neurons that interact with body wall muscles posterior to the head and neck.

### 4.2 Sensory neuron

Being the only neuron in the proposed circuit that receives input from the external environment rather than other neurons, the input term I_AWA_ is given by the equation

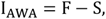

where F represents the fast component of the sensory response and S the slow component (17). This model of AWA activation allows the response of this neuron to recapitulate the experimentally observed response of AWA membrane potential to chemical stimulus during natural turning behavior (37). The fast and slow sensory response is given as follows:

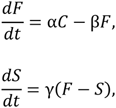

where α is the depolarization rate, β is the leak rate, γ is the repolarization rate, and *C = C*(*x,y*) is the strength of the food gradient at coordinates (*x*, *y*).

### 4.3 Muscles and stretch receptors

Head and VNC motor neurons, body wall muscles, and ventral nerve cord circuitry were modeled based on (16). Briefly, 24 dorsal and 24 ventral muscles were modeled as damped springs with activation *A^k^*_M,*m*_:

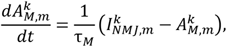

where *I^k^*_NMJ,*m*_ is the current driving the muscles. Head and neck muscles (*m* = [1,6]) are driven by SMD and RMD motor neuronal activity according to:

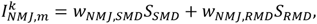

where *k =* {*D*, *V*} and *S* denotes the output (*y*, see 4.4) of a given neuron.

Downstream muscles in the body (*m* = [7, 24]) are driven by ventral nerve cord circuit D- and B-class motor neurons according to:

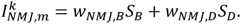

Both SMD-class motor neurons in the head circuit and B-class motor neurons in the ventral nerve cord circuit receive stretch receptor input. SMD-class motor neuron stretch receptor current is summed over contributions from 14 mechanical elements associated with (*m* = [4, 9]), comprising neck muscles and muscles associated with the first ventral nerve cord unit. B-class motor neuron stretch receptor current is summed over contributions from the six mechanical elements anterior to the anterior-most muscle that neuron innervates. Stretch receptor activation is modelled as a weighted linear function of muscle length (29):

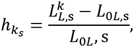

where *L_OL,s_* is the segment rest length and L^k^_L,s_ is the current length of the *k*th side (dorsal/ventral) of the *s*th segment.

### 4.4 Neural model

Neurons were modelled based on prior established work (16, 27, 47):

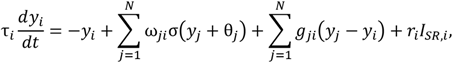

where *y* is the neuronal activation, τ is the time constant, and ω_s_ is the value of the self-connection. Graded synaptic output was modelled as a sigmoidal function σ(*x*) of presynaptic activation:

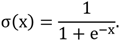

Gap junction conductance *g_ji_* is modeled as bidirectional ohmic resistances. Stretch receptor input *I_SR,i_* is computed as the sum of mechanical elements contributing to that stretch receptor.

### 4.5 Numerical methods

The model was implemented in C++ and Python. Muscle output was solved with explicit Euler integration with a step of 1ms, and neuronal output was solved with 4^th^-order Runge-Kutta integration to minimize numerical instability in these approximations.

### 4.6 Food gradient

The chemical attractant concentration *c(x, y)* was modelled using a conical gradient from a point source in which the concentration is proportional to the Euclidean distance between the tip of the model worm’s head to the gradient peak (29):

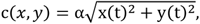

where *x(t)* and *y(t)* indicate the Euclidean distance to the gradient peak and the scalar α determines the steepness of the gradient. Gaussian gradients were modelled as:

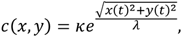

where κ denotes the amplitude of the gradient and λ denotes the diffusion coefficient.

### 4.7 Parameter optimization

Overall, the model has 47 free parameters, consisting of three chemosensory input parameters (α, β, γ); seven time constants (τ) and seven biases (θ), one for each neuronal class; six self-connections, one for each neuronal class except AWA (which solely receives input from the external environment); four neuromuscular junctions, one for each motor neuron class; two stretch-receptor gains for SMD and B stretch receptors; weights (ω) in the head neuronal circuit for (a) nine chemical synapses (i) from AWA to RIM, (ii) from RIM to RIA, (iii) from RIA to RMD, (iv) from RIA to SMD, (v) from RMD to RIA, (vi) from SMD to RIA, (vii) between SMD, (viii) from SMD to RMD, and (ix) between RMD, and (b) two electrical synapses between (i) RMD and (ii) SMD and RMD; weights in the VNC neural unit for (a) three chemical synaptic connections within a given unit (i) from B- to D- motorneurons ipsilaterally, (ii) from B- to D- motor neurons contralaterally, and (iii) between D-motor neurons, one intra-unit electrical synapse between D-motor neurons, and three electrical synapses across units between (i) neighboring D-motor neurons, (ii) neighboring B-motor neurons, and (iii) contralateral neighboring B-motor neurons. Values for all these parameters were optimized using a genetic/evolutionary algorithm (cite) implemented in C++ using the JSON for Modern C++ library (Lohmann, N. (2025). JSON for Modern C++ (Version 3.12.0).

For parameter optimization, an initial population of 100 worms (parameter sets) was generated by randomizing each parameter within a given range. Adapted from prior work (27), fitness was represented as the chemotaxis index (CI), given as:

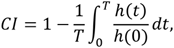

where *h(t)* is the Euclidean distance to the food source at time *t, h(0)* is the model worm’s initial distance from the food source (0.4 cm), and *T* is the total simulated assay time (100 s). For simplicity, negative *CI* values were set to zero. The fitness of an individual worm was defined as the average *CI* over five assays, where food was placed in a different location and gradient strength α was randomized within the range [-1.0, -0.1]. For each generation, individuals were selected based on elitism and fitness proportional selection, underwent one-point crossover at a rate of 0.7 and mutation at a rate of 0.005, and were then evaluated for the subsequent generation for a total of 100 generations.

### 4.8 Terminology

All analysis is based on the direction of translation, defined as any two points separated by one cycle of locomotion, i.e., 2π. A cycle of locomotion consists of one dorsal and one ventral turn. Dorsal and ventral turns are defined as when the turning angle φ is greater than or less than zero, respectively. Turning bias is defined as the sum of the turning angle φ over one cycle of locomotion. Bearing is defined as the angle between the line of steepest ascent in the attractant concentration gradient field and the direction of translation.

For each video, food is placed 8 mm from worm’s initial position and the simulation runs for 100 seconds with a 0.01 time step. Cardinal directions are used to indicate the angle difference between the food source and the worm’s initial heading angle: *north* indicates a 90° difference (**#Video 1**), *south* indicates 270° (**#Video 2**), *east* indicates 0° (**#Video 3**), and *west* indicates a 180° difference (**#Video 4**).

### 4.9 Statistical analysis

All values listed as (µ ± σ*_x_*) indicate mean ± S.E.M. Welch’s t-tests are described by giving the degrees of freedom, t-statistic value, and p-value. Correlation coefficients (r) were calculated using the Pearson correlation method. One-way ANOVA is reported with degrees of freedom, F-statistic, and p-value. Post-hoc Tukey HSD was used to define statistically similar groups following ANOVA. Area under the curve (AUC) was calculated across 30 seconds and reported with mean ± S.E.M.

## Acknowledgements

The authors would like to thank Michael Ivanitskiy for help with initial coding of the model, and Zihan Zhou and Conghao Jin for initial parameter explorations.

## 6. Data availability

Data related to this work are provided in the Supplementary Figures and Videos. The code for the model and genetic algorithm can be found at: https://github.com/adasquires/celegans-chemotaxis.

---

**Supplementary Figure 1.**
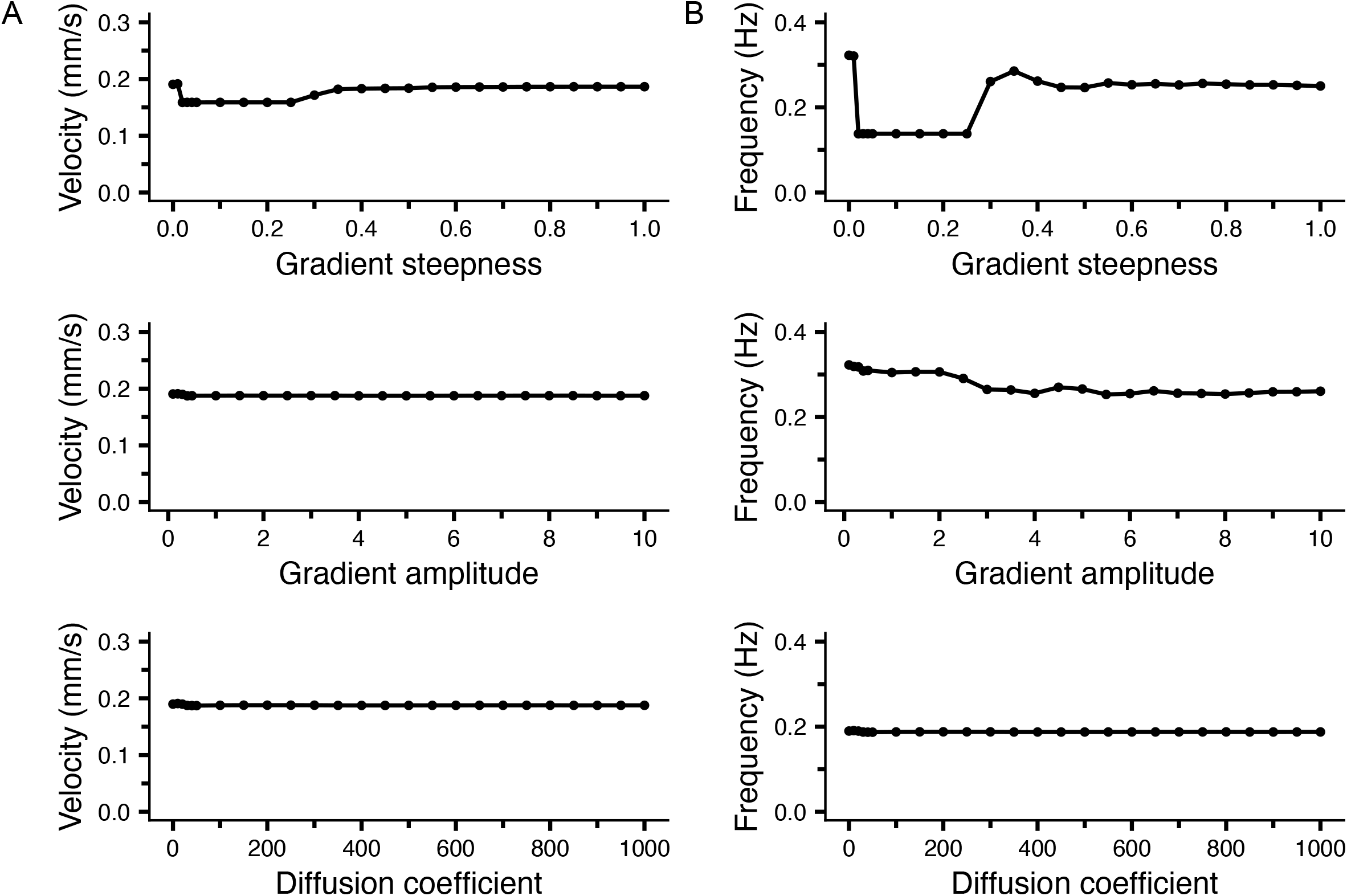
Locomotion analysis. Analysis of model worm’s (A) average velocity in mm/s and (B) frequency in Hz across a simulation in attractant concentration gradient fields with variable conical gradient steepness (top), Gaussian gradient amplitudes (middle), and Gaussian diffusion coefficients (bottom). Data is averaged from four simulations with different food positions, placed 10mm from the worm’s initial position with an angle difference of 0°, 90°, 180°, and 270° from the worm’s initial heading angle.

**Supplementary Figure 2.**
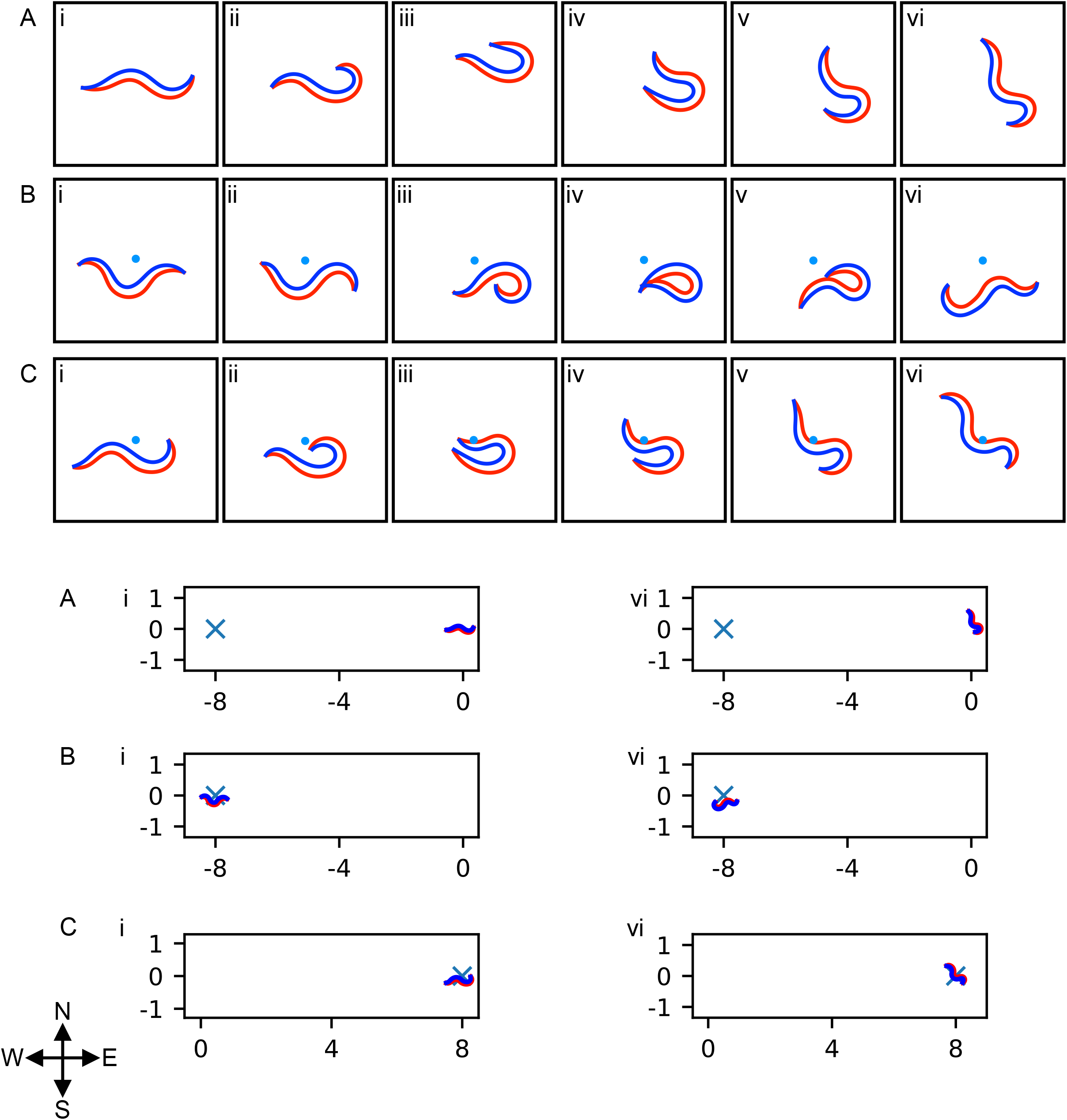
Model worms’ body shape during sharp/omega-like turns in simulations of the optimized model. Top three rows of panels: Sequential snapshots of (A) initial sharp turn when the food source is placed “west” of the worm, (B) sharp turn at the end of “west” simulation to remain in proximity of the food source (blue dot), and (C) sharp turn at the end of “east” simulation to remain in proximity of the food source (blue dot). In all (i) panels the initial heading of the worm is to the “east”. Bottom set of panels: First (left panels) and last (right panels) snapshots of the A, B, and C simulations zoomed out, to show where the worm is in relation to the food source, which is marked with an x symbol. Dorsal side of body is depicted in red and ventral side of body is depicted in blue. See Methods 4.7 for terminology.

**Supplementary Figure 3.**
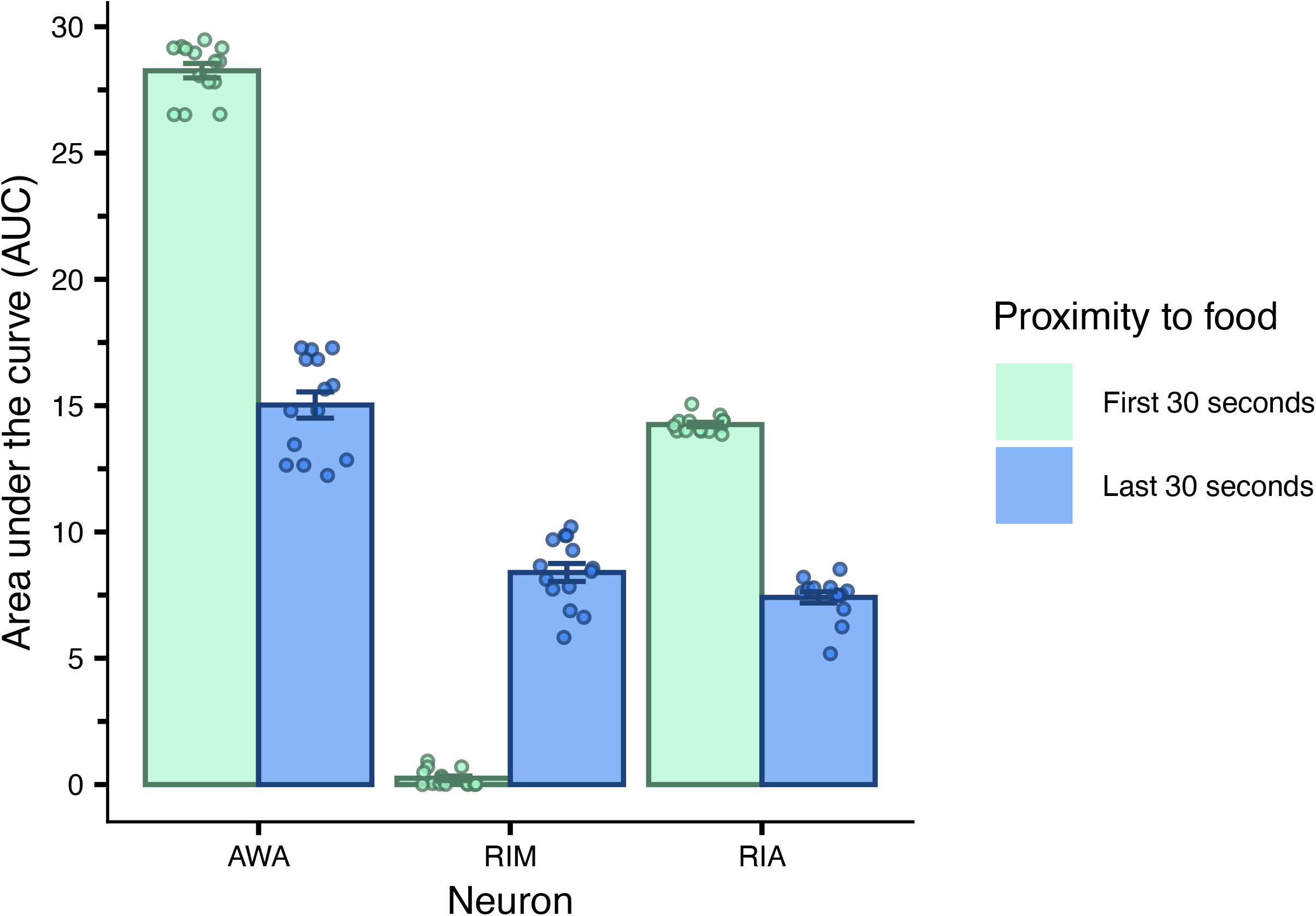
Area under the curve (AUC) analysis. AUC of AWA, RIM, and RIA neuronal activity across 30 seconds at the beginning of a simulation (magenta), when the worm is distal to the food source and conducting shallow turns, and at the end of a simulation (blue), when the worm is proximal to the food source and conducting sharp turns. AUC was performed across four simulations with variable food positions and averaged on the bar plot.

**Supplementary Figure 4.**
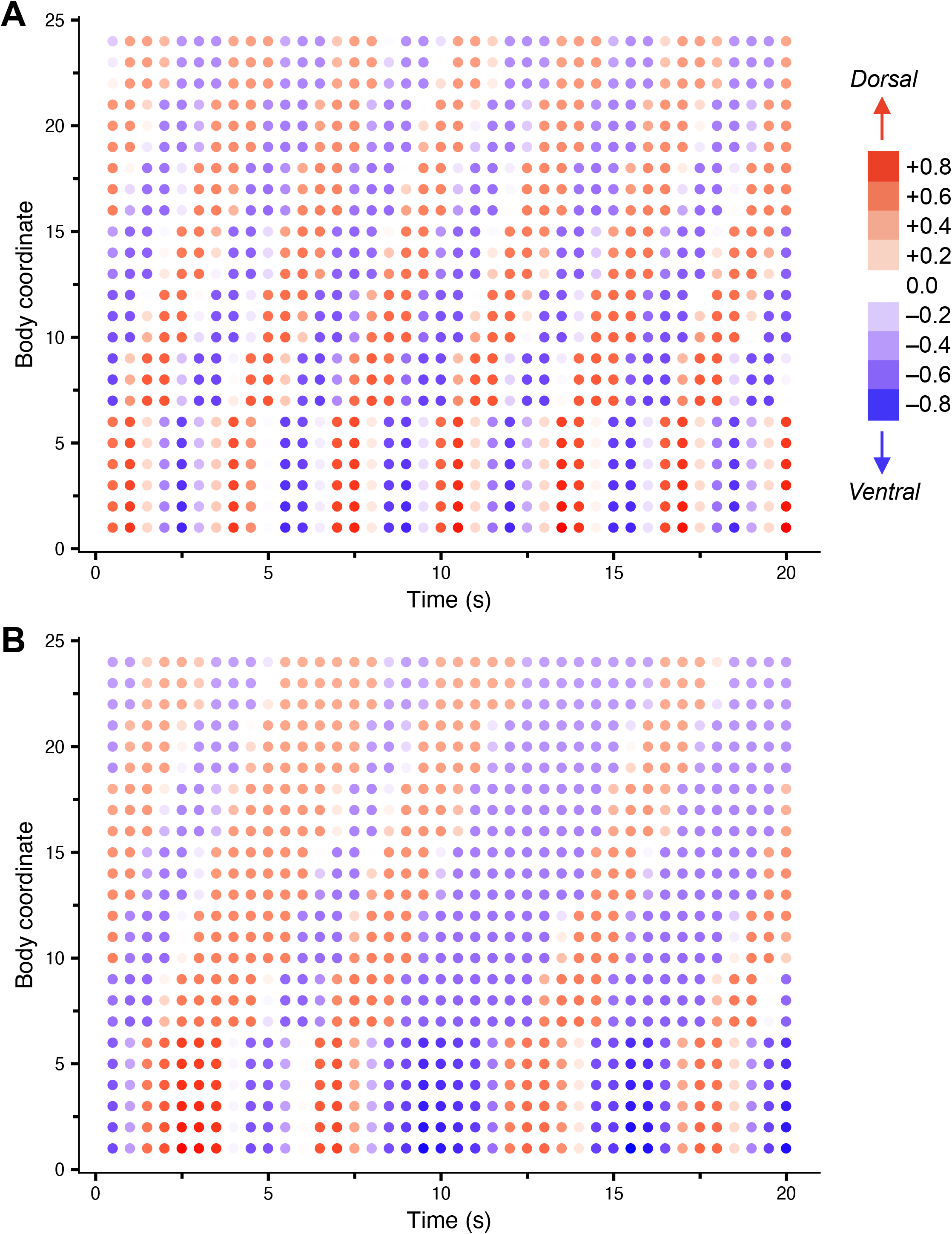
Muscle activity propagates oscillations through the body to support sharp turning in chemotaxis. Kymographs of muscle activity in (A) shallow turning and (B) sharp turning. All muscle activity is recorded as a positive number between [0.0, 1.0]. At each time point (x-axis) and body coordinate (y-axis) ventral muscle activation is subtracted from dorsal muscle activation. Positive values indicate dorsal muscle activation bias (red) and negative values indicate ventral muscle activation bias (blue). Points are calculated at each time point across 20 seconds and for each of the 24 body segments of the model, with body segment 1 representing the most anterior muscles in the head and body segment 24 representing the most posterior muscles in the tail of the model. Both plots are generated from the same 100 second simulation: (A) shows an early time interval in the simulation when the worm is distal to the food source and completing shallow turns (between [10, 30] seconds within a 100 second simulation), (B) shows a late time interval in the simulation when the worm is proximal to the food source and completing sharp turns (between [50, 70] seconds within a 100 second simulation).

**Supplementary Figure 5.**
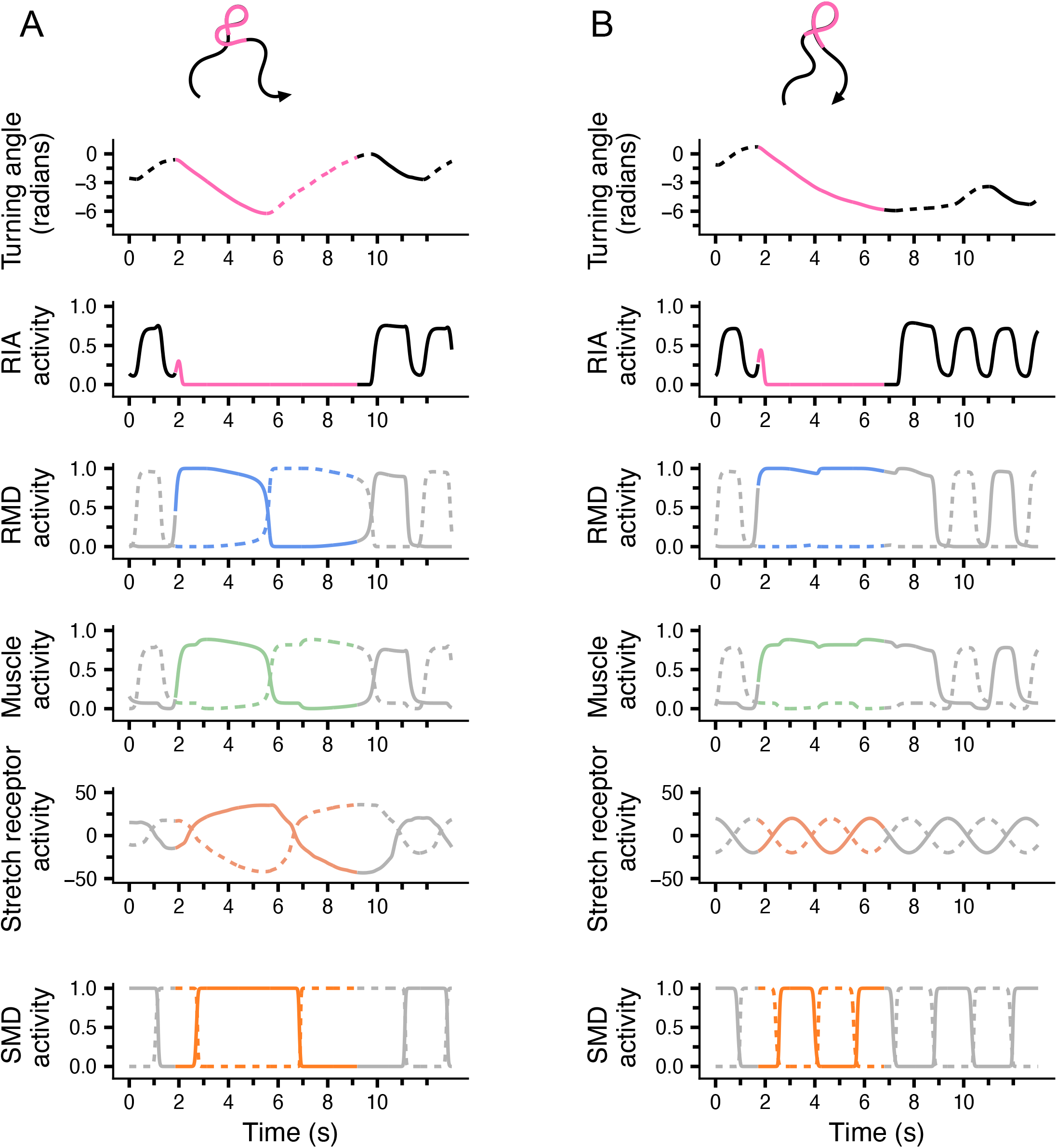
Analysis of proprioceptive feedback in chemotaxis. Comparison of turning angle, motor neuron, muscle, and stretch receptor response to a down-step in concentration, when stretch receptor activity is either (A) allowed to respond to sharp turns or (B) manipulated to oscillate only in time with shallow turning. Colored lines represent the time points in which the concentration step occurs. Top: representative behavioral trajectory across the simulation. In (A), stretch receptor response allows the worm to perform a sharp turn in both directions consecutively, forming a “figure-eight” pattern. In (B), lack of stretch receptor response results in the worm completing a sharp turn in one direction and then stalling before concentration steps back up to 0.0, whereupon it begins completing shallow turns. This stalling is represented in the behavioral trace as a lack of a “figure-eight” pattern that characterizes consecutive sharp turns.

**Supplementary Figure 6.**
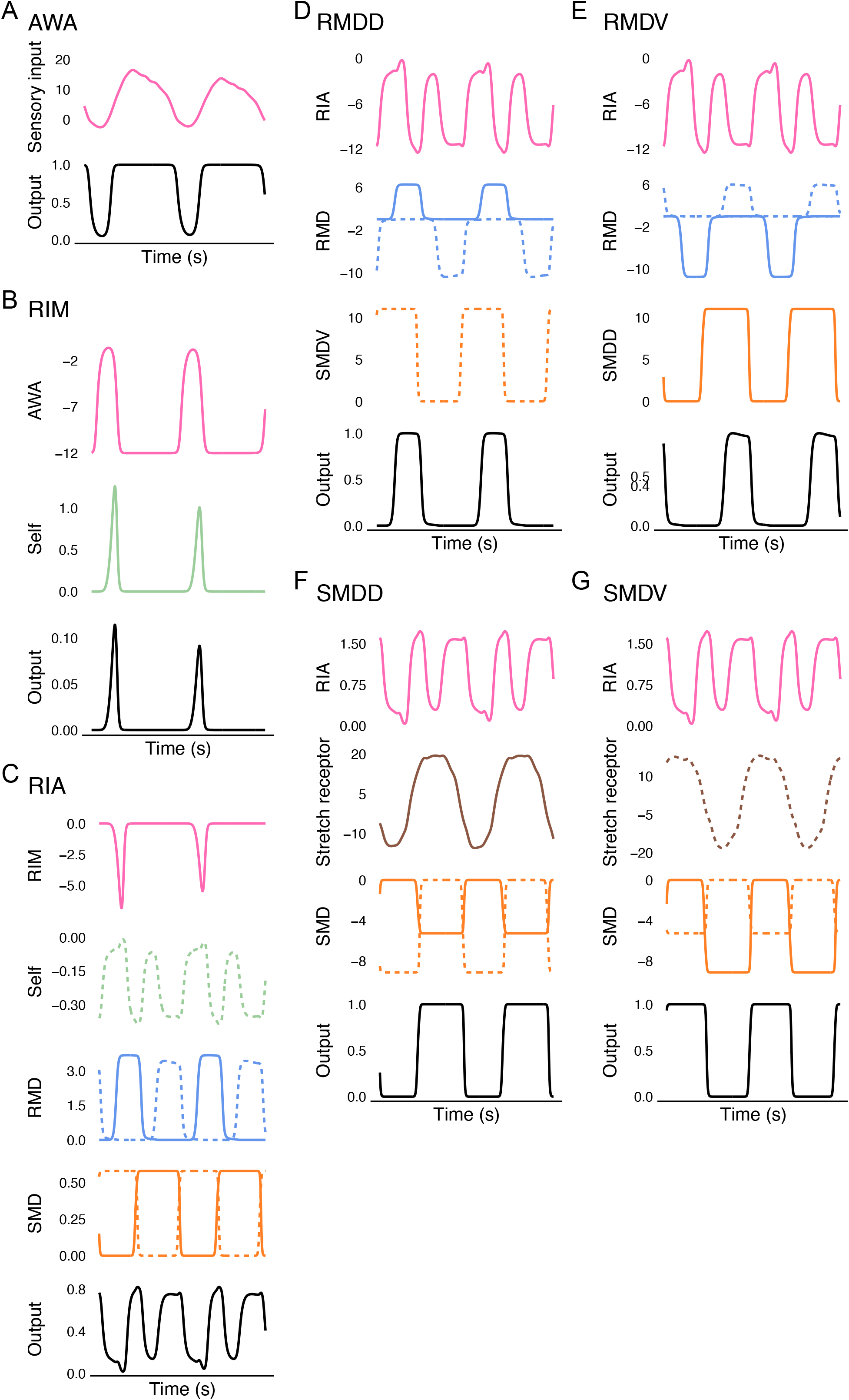
Tolerance to noise. Gaussian noise with a mean of 0.3 and an amplitude of 3.0 is applied individually to each head neuron. Chemotaxis index is recorded across ten simulations with the same initial heading angle and variable food locations. Individual points represent values from individual simulations. Bar plots represent the average chemotaxis index of each manipulation across simulations. Dashed line indicates chemotaxis index of deterministic model.

**Supplementary Figure 7.**
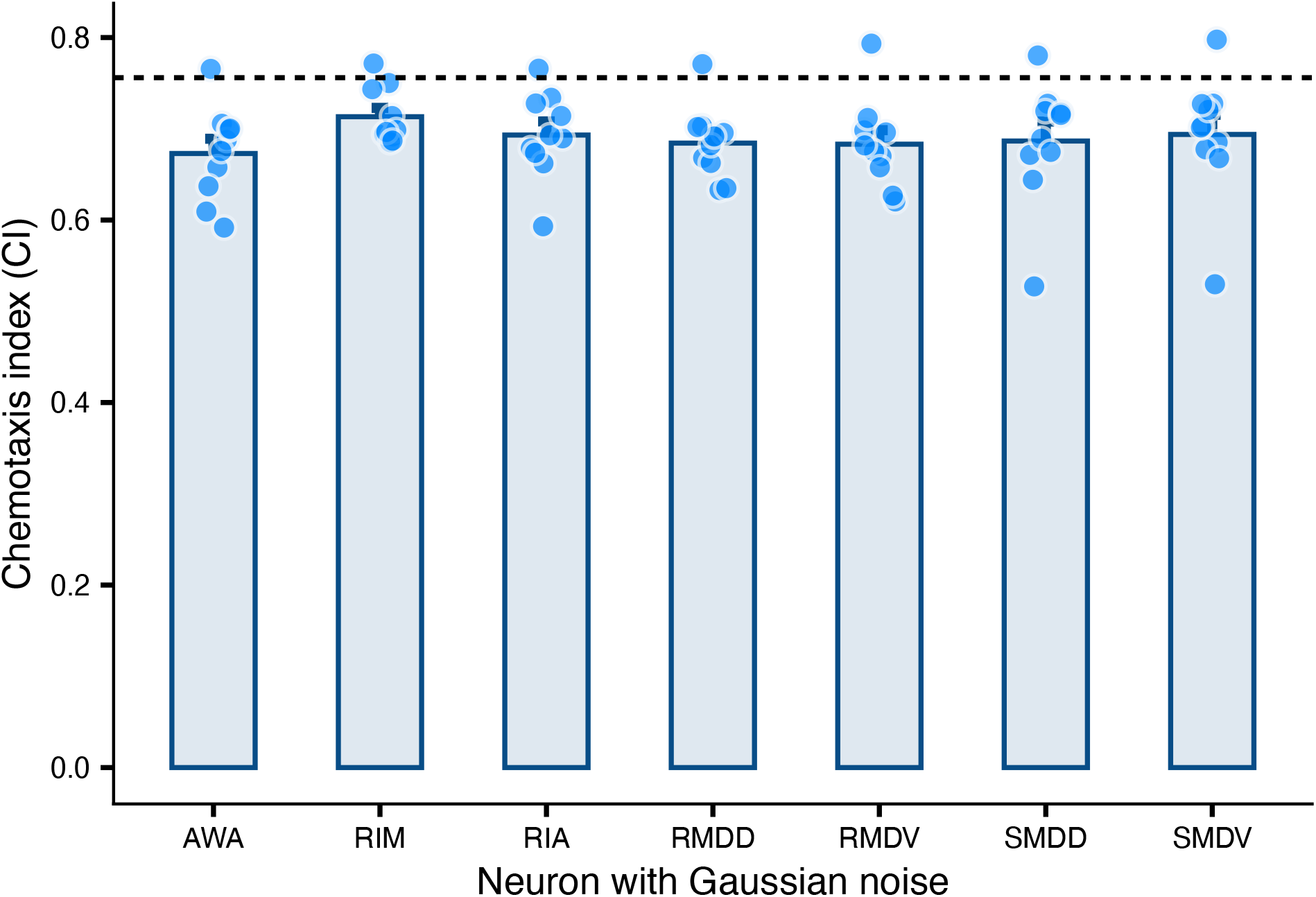
Input-output relationships of head neurons. In all graphs, input dorsal neurons (RMDD, SMDD) are indicated in solid lines, and input from ventral neurons (RMDV, SMDV) are indicated in dashed lines. (A) AWA sensory input (pink) and output (black). (B) RIM input from AWA (pink), self-connection (green), and output (black). (C) RIA input from RIM (pink), self-connection (green), RMD (blue), SMD (orange), and output (black). (D) RMDD input from RIA (pink), RMD (blue, solid line is a self-connection), SMDV (orange), and output (black). (E) RMDV input from RIA (pink), RMD (blue, dashed line is self- connection), SMDD (orange) and output (black). (F) SMDD input from RIA (pink), dorsal stretch receptor (brown), SMD (orange, solid line is a self-connection), and output (black). (G) SMDV input from RIA (pink), stretch receptors (brown), SMD (orange, dashed line is a self- connection), and output (black).

**Supplementary Figure 8.**
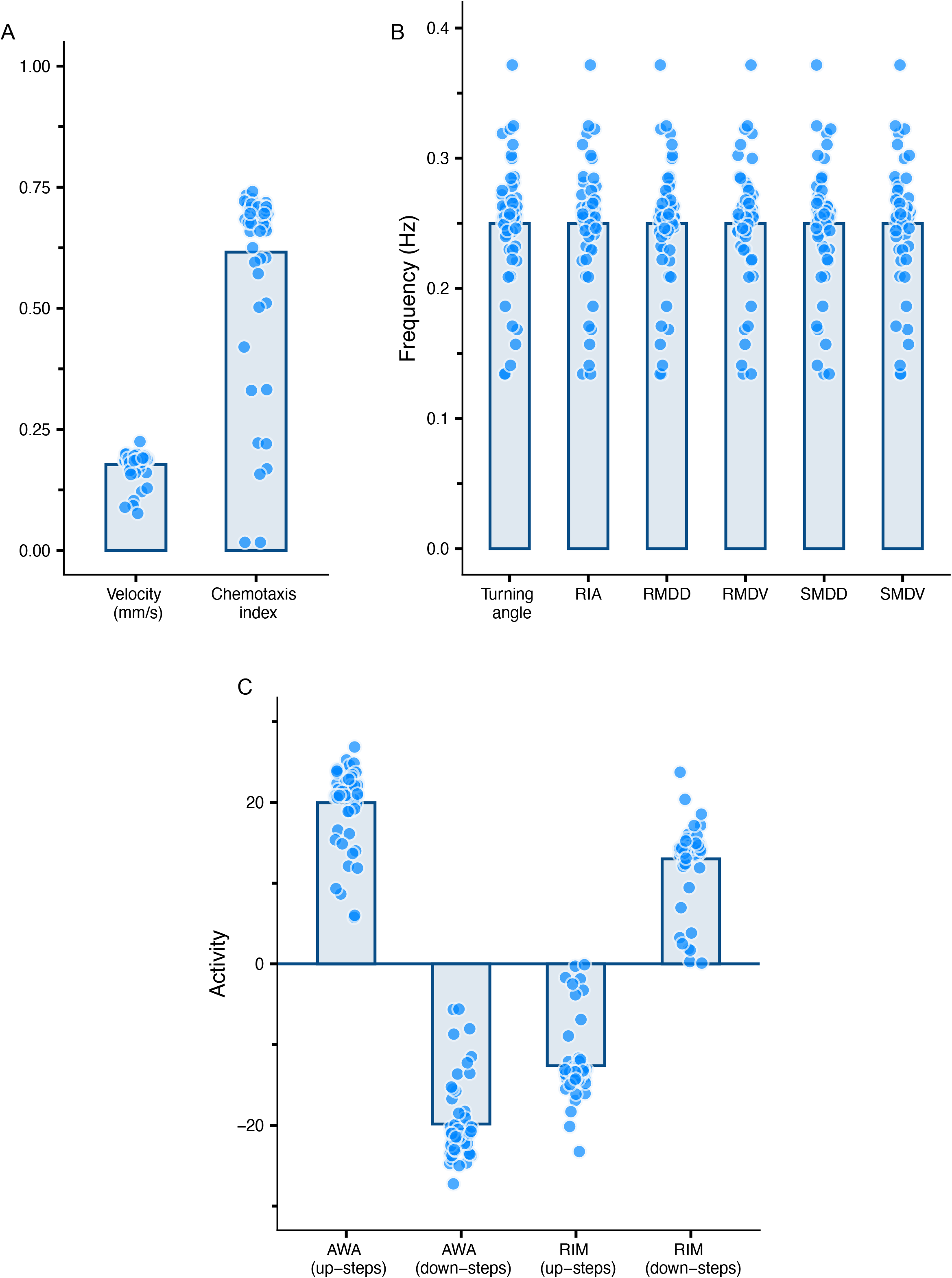
Analysis of family of optimization-derived model parameter sets. Analysis was conducted on sensitivity of model results for parameter sets within the ranges provided in Table 1. Each individual point represents the averaged value across four simulations with variable food locations when one parameter was altered to the extreme value of its range. Bar plots show averaged values. All values are gathered from simulations of 100 seconds with food placed 8 mm from the worm’s initial position. (A) Velocity (mm/s) and chemotaxis index. (B) Frequency (Hz) of turning angle, RIA, RMD, and SMD neuronal activity. (C) Cumulative activity of AWA and RIM in up-steps (when concentration is increasing) and down-steps (when concentration is decreasing). Cumulative activity is defined here as the summed change in neuronal output. Across a 100 second simulation in which the worm spends approximately equal amounts of time experiencing increases and decreases in concentration, the summed changed in AWA and RIM activity was recorded across all responses to concentration increases and all responses to concentration decreases.

**Video 1.** Animation of simulation with food placed north of worm (see Methods 4.7 for terminology).

**Video 2.** Animation of simulation with food placed south of worm (see Methods 4.7 for terminology).

**Video 3.** Animation of simulation with food placed east of worm (see Methods 4.7 for terminology).

**Video 4.** Animation of simulation with food placed west of worm (see Methods 4.7 for terminology).

